# Gene expression and RNA splicing explain large proportions of the heritability for complex traits in cattle

**DOI:** 10.1101/2022.05.30.494093

**Authors:** Ruidong Xiang, Lingzhao Fang, Shuli Liu, Iona M. Macleod, Zhiqian Liu, Edmond J. Breen, Yahui Gao, George E. Liu, Albert Tenesa, CattleGTEx Consortium, Brett A Mason, Amanda J. Chamberlain, Naomi R. Wray, Michael E. Goddard

## Abstract

Many quantitative trait loci (QTL) are located in non-coding genomic regions. Therefore, QTL are assumed to affect gene regulation. Gene expression and RNA splicing are primary steps of transcription so QTL changing gene expression (eQTL) or RNA splicing (sQTL) are expected to significantly contribute to phenotypic variations. Here, we quantify the contribution of eQTL and sQTL detected from 16 tissues (N~5,000) to 37 complex traits of ~120k cattle. Using Bayesian methods, we show that including more regulatory variants in the model explains larger proportions of heritability. Across traits, *cis* and *trans* eQTL and sQTL detected from 16 tissues jointly explain ~70% (SE=0.5%) of heritability, 44% more than expected from the same number of random variants, where trans e/sQTL contribute 24% (14% more than expected). Multi-tissue *cis* and *trans* e/sQTL also explain 71% (SE=0.3%) of heritability for the metabolome, demonstrating the essential role of proximal and distal regulatory variants in shaping mammalian phenotypes.

## Introduction

Understanding how DNA variants shape phenotype is a central goal in genetics and biology. Most mammalian phenotypes are influenced by the accumulated effects of many quantitative trait loci (QTL) from non-coding regions. Since non-coding regions are usually involved in gene regulation, numerous human studies have mapped regulatory loci, including QTL affecting gene expression (eQTL) ^1,2^ and RNA splicing (sQTL) ^3^, in the expectation that they would explain variation in complex traits.

Significant efforts in mapping regulatory variants in other species have also been initiated very recently, including large animal species. A Cattle Genotype-Tissue Expression (CattleGTEx) ^4^ consortium, as part of Farm animal GTEx (FarmGTEx), has been launched along with new priorities for the Functional Annotation of ANimal Genomes (FAANG) ^5,6^ consortium. Genome-wide association studies (GWAS) of cattle are now carried out in more than 100,000 individuals ^7,8^ to identify causal QTL for dozens of traits. Therefore, there are unique opportunities in non-human species to dissect the impact of regulatory variants on mammalian complex traits.

Despite being biologically important, regulatory variants have been reported to contribute only a small part to variation in mammalian complex traits. For example, a recent human study suggested that around 11% of trait heritability is attributable to eQTL ^9^. In the evaluation of published human data, Connally et al ^10^ proposed the term ‘missing regulation’ to describe the discordance between eQTL and QTL. In cattle, limited overlaps between eQTL and QTL has been reported ^11^ and the total contribution of eQTL to the heritability of cattle traits was also around 10% ^12^. In human studies investigating the contributions to trait variation eQTL and sQTL have not, to date, been analysed together.

Herein, we address the contribution of regulatory variants to mammalian complex traits with a comprehensive analysis of cattle data. We mapped eQTL and sQTL from transcriptomes across 16 tissues in more than 40 breeds from ~5,000 cattle. In another ~120k Australian cattle, we use a Bayesian mixture model allowing prior information that a variant affects gene regulation ^13^, to estimate the genetic variance explained across 37 traits by *cis* and *trans* eQTL and sQTL in single tissues as well as over multiple tissues. We replicate the analysis in metabolomic traits assayed by liquid chromatography-mass spectrometry and found that regulatory variants explain a large proportion of genetic variance.

## Results

Expression QTL and sQTL were mapped in 16 tissues in either newly generated data or data obtained from CattleGTEx V0 ^4^ in tissues with sample size > 100 (Supplementary Table 1) using a linear mixed model approach ^14^ (see Methods). There were 4,725 independent samples across 16 tissues and on average 295 samples per tissue. We mapped *cis* (±1Mb of gene or intron) e/sQTL with p < 5 × 10^-6^ in the association mapping. More stringent criteria were applied to the selection of *trans* e/sQTL (from different chromosomes to the gene or intron, see Methods). e/sQTL were also categorised as within or outside regions under Chromatin Immunoprecipitation sequencing (ChIP-seq) peaks ^6^ (see Methods). A metaanalyses across 16 tissues was conducted to identify multi-tissue eQTL and sQTL (see Methods). We then classified >1.8 million LD-pruned (*r*^2^ < 0.9) genome-wide variants in ~120k Australian cattle into 13 classes based on whether they were eQTL, sQTL, or both eQTL and sQTL (esQTL) that act in *cis* or *trans* and whether they were *cis/trans* e/sQTL under ChIP-seq peaks at both single-tissue and multi-tissue levels (Supplementary Table 2).

Each of the 37 complex traits (Supplementary Table 3) was analysed with the BayesRC method ^13^. Like BayesR ^15,16^, BayesRC assumes that the effect of a variant on a complex trait is drawn from a mixture of 4 normal distributions with mean=0 and variances of zero, 0.0001, 0.001, or 0.01 times the genetic variance. However, in Bayes RC, the variants are placed into classes based on prior information (e.g., with regulatory evidence), and the proportion of each distribution in the mixture is allowed to vary between classes. This allowed us to quantify the relative proportion of heritability attributable to classes of *cis* and *trans* eQTL and sQTL. Here the heritability is based on additive genetic variance due to sequence variants, equivalent to the term “SNP-heritability” in human genetics. In BayesRC, the proportion of heritability explained was also estimated for a class of ‘remaining variants’ with no regulatory evidence. We used this estimate together with its ratio of genomic size to other classes of regulatory variants to derive an expected proportion of heritability explained by each class of variants assuming they explained the same amount per variant as the remaining class (Table 1 and Methods).

**Table 1.** Summary of the proportion (%) of heritability and trait-associated variants (QTL) in expression quantitative trait loci (eQTL), RNA splicing QTL (sQTL), variants that are both eQTL and sQTL (esQTL) and those e/sQTL under histone marks (“_Histone”). Within each class, the total number of variants (N class) and their genome proportion (% class), the number of variants with small effects, medium effects and large effects averaged across 16 tissues and 37 traits are given. These numbers are used to estimate the observed heritability explained (O[% *h^2^*]) and the proportion of QTL in each class (O[% QTL]). The number of variants within the remaining class (no regulatory evidence) is used to estimate the expected proportion of heritability explained (E[% *h^2^*]) and proportion of QTL in each class (E[% QTL]).

| Tissue | Class | N class | % class | Small | Medium | Large | $O[\% h^2]$ (se) | $E[\% h^2]$ (se) | $O[\% \text{ QTL}]$ (se) | $E[\% \text{ QTL}]$ (se) |
| --- | --- | --- | --- | --- | --- | --- | --- | --- | --- | --- |
| Single-tissue | cis.eQTL_Histone | 3011 | 0.16 | 73.4(2) | 7.6(0.2) | 0.3(0.0) | 1.70(0.04) | 0.12(0.00) | 5.19(0.18) | 0.001(0.000) |
|  | cis.eQTL | 4910 | 0.26 | 93.1(3) | 9.2(0.3) | 0.3(0.0) | 2.08(0.05) | 0.20(0.01) | 3.93(0.14) | 0.001(0.000) |
|  | cis.sQTL_Histone | 9835 | 0.52 | 127.7(4) | 11.1(0.3) | 0.4(0.0) | 2.66(0.07) | 0.40(0.02) | 2.52(0.12) | 0.002(0.000) |
|  | cis.sQTL | 16386 | 0.87 | 197.2(8) | 13.9(0.5) | 0.4(0.0) | 3.57(0.12) | 0.66(0.03) | 2.07(0.09) | 0.003(0.000) |
|  | cis.esQTL_Histone | 1928 | 0.10 | 50.0(2) | 5.4(0.2) | 0.3(0.0) | 1.29(0.04) | 0.08(0.01) | 11.20(0.54) | 0.000(0.000) |
|  | cis.esQTL | 2669 | 0.14 | 56.4(2) | 5.8(0.2) | 0.3(0.0) | 1.38(0.04) | 0.11(0.01) | 9.66(0.53) | 0.001(0.000) |
|  | trans.eQTL_Histone | 939 | 0.05 | 53.9(1) | 5.8(0.2) | 0.2(0.0) | 1.30(0.03) | 0.04(0.00) | 11.46(0.45) | 0.000(0.000) |
|  | trans.eQTL | 2064 | 0.11 | 74.5(2) | 8.1(0.3) | 0.3(0.0) | 1.74(0.04) | 0.09(0.00) | 7.57(0.33) | 0.000(0.000) |
|  | trans.sQTL_Histone | 9891 | 0.53 | 115.7(2) | 12.3(0.4) | 0.3(0.0) | 2.60(0.06) | 0.41(0.01) | 1.60(0.06) | 0.002(0.000) |
|  | trans.sQTL | 22191 | 1.18 | 181.2(3) | 16.6(0.5) | 0.4(0.0) | 3.67(0.07) | 0.93(0.01) | 1.01(0.02) | 0.005(0.000) |
|  | trans.esQTL_Histone | 847 | 0.04 | 46.8(1) | 5.2(0.2) | 0.2(0.0) | 1.16(0.03) | 0.04(0.00) | 10.13(0.50) | 0.000(0.000) |
|  | trans.esQTL | 1893 | 0.10 | 63.1(2) | 6.8(0.2) | 0.3(0.0) | 1.49(0.04) | 0.08(0.00) | 7.12(0.46) | 0.000(0.000) |
|  | remaining | 1805933 | 95.9 | 6726.8(54) | 98.1(2.1) | 1.7(0.1) | 75.38(0.41) | 75.38(0.41) | 0.38(0.00) | 0.38(0.00) |
| Multi-tissue | cis.eQTL_Histone | 343 | 0.02 | 33.1(4) | 4.0(0.4) | 0.2(0.0) | 0.87(0.08) | 0.01(0.00) | 10.88(1.12) | 0.000(0.000) |
|  | cis.eQTL | 1576 | 0.08 | 51.8(6) | 5.5(0.6) | 0.2(0.0) | 1.22(0.11) | 0.05(0.00) | 3.65(0.42) | 0.000(0.000) |
|  | cis.sQTL_Histone | 67720 | 3.60 | 385.5(20) | 22.8(2.3) | 0.5(0.1) | 6.30(0.26) | 2.03(0.10) | 0.60(0.03) | 0.018(0.001) |
|  | cis.sQTL | 184798 | 9.82 | 1225.7(62) | 32.8(3.9) | 0.5(0.0) | 15.16(0.66) | 5.54(0.29) | 0.68(0.03) | 0.048(0.002) |
|  | cis.esQTL_Histone | 105956 | 5.63 | 557.2(27) | 23.8(2.4) | 0.7(0.1) | 8.26(0.38) | 3.18(0.16) | 0.55(0.02) | 0.028(0.001) |
|  | cis.esQTL | 169434 | 9.00 | 1036.6(47) | 29.7(3.1) | 0.8(0.2) | 13.50(0.77) | 5.08(0.26) | 0.63(0.03) | 0.044(0.002) |
|  | trans.eQTL_Histone | 53187 | 2.83 | 243.9(13) | 14.9(1.9) | 0.4(0.0) | 4.09(0.19) | 1.59(0.08) | 0.49(0.02) | 0.014(0.001) |
|  | trans.eQTL | 173939 | 9.24 | 743.9(33) | 17.4(1.6) | 0.4(0.0) | 9.04(0.29) | 5.22(0.27) | 0.44(0.02) | 0.045(0.002) |
|  | trans.sQTL_Histone | 7076 | 0.38 | 106.8(10) | 8.9(0.9) | 0.3(0.0) | 2.10(0.16) | 0.21(0.01) | 1.64(0.14) | 0.002(0.000) |
|  | trans.sQTL | 40618 | 2.16 | 258.5(22) | 14.2(1.3) | 0.4(0.0) | 4.11(0.26) | 1.22(0.06) | 0.67(0.05) | 0.011(0.001) |
|  | trans.esQTL_Histone | 5603 | 0.30 | 76.2(7) | 6.8(0.5) | 0.3(0.0) | 1.60(0.11) | 0.17(0.01) | 1.49(0.14) | 0.001(0.000) |
|  | trans.esQTL | 44089 | 2.34 | 168.4(14) | 11.0(1.2) | 0.3(0.0) | 2.91(0.20) | 1.32(0.07) | 0.41(0.03) | 0.011(0.001) |
|  | remaining | 1028161 | 54.6 | 2708.2(138) | 44.1(9.5) | 0.9(0.2) | 30.83(1.59) | 30.83(1.59) | 0.27(0.01) | 0.27(0.01) |

The BayesRC analysis was performed when the regulatory classes were defined based on each tissue one at a time (e.g., eQTL called from single-tissue) and when the classes were defined based on all tissues together (e.g., eQTL called from 16 tissues). Table 1 gives the BayesRC results for the 13 prior classes (fitted jointly) averaged across the 37 traits for both the single-tissue analyses and the multiple-tissue analysis (the single-tissue results are averaged across all single-tissue analyses). In single-tissue and multiple-tissue analyses, for all 12 regulatory classes, the proportion of variants that had an effect on phenotypes (small, medium, or large) was greater than for the remaining class that were not regulatory variants. Also, within the variants that affected phenotype, the proportion of variants with medium or large effect was greater for the 12 regulatory classes than for the remaining class. Consequently, the variance explained by the 12 regulatory classes was higher than expected if they explained the same amount per variant as the remaining classes.

In the multiple tissue analysis, when all tissues were used to define regulatory variants, more variants were classified as regulatory and fewer variants remain in the ‘remaining’ class. Also, within the regulatory classes, more variants affected both gene expression and RNA splicing and so the esQTL class had more variants and the eQTL and sQTL classes had fewer variants (Table 1). As a result of the larger number of regulatory variants discovered, the multiple-tissue analysis found that 70% of the genetic variance was explained by regulatory variants (44% more than expected by the same number of random variants) whereas the average of the single-tissue analyses was 25% (22% more than expected). As the multi-tissue analysis had more than a 16-fold increase in sample size compared to each single-tissue analysis (4725 VS 295), our results suggest that the higher information content in the multitissue e/sQTL was likely due to the greater power in their mapping.

Including more regulatory variants in the model increased the heritability explained by them, with the largest proportions of heritability explained by eQTL and sQTL detected from all tissues analysed jointly (Figure 1a-c). To further illustrate this, as well as the 13 classes defined by both eQTL and sQTL, we performed BayesRC analyses using 5 classes defined only by eQTL or sQTL (Supplementary Table 4). Based on e/sQTL detected from single tissues averaged across tissues and traits, when eQTL and sQTL were analysed separately, *cis* and *trans* eQTL explained 5.6% (SE=0.1%) and 4.1% (0.1%) of heritability respectively, and *cis* and *trans* sQTL explained 8.3% (0.2%) and 7.6% (0.1%) of heritability respectively (Supplementary Table 4). When eQTL and sQTL were analysed jointly in the single-tissue scenarios, *cis* and *trans* esQTL explained 12.6% (0.1%,) and 11.9% (0.1%) of heritability respectively (Table 1). Based on e/sQTL detected from multiple tissues across traits, when eQTL and sQTL were analysed separately, *cis* and *trans* eQTL explained 29% (1%) and 21% (0.5%) of heritability respectively, and *cis* and *trans* sQTL explained 58% (1%) and 8% (0.4%) of heritability respectively (Supplementary Table 4). When eQTL and sQTL were analysed jointly in the multi-tissue scenarios, *cis* and *trans* esQTL explained 45% (0.5%) and 24% (0.3%) of heritability respectively (Table 1). A full list of partitioned heritability across tissues and traits can be found in Supplementary Data 1.

**Figure 1.**
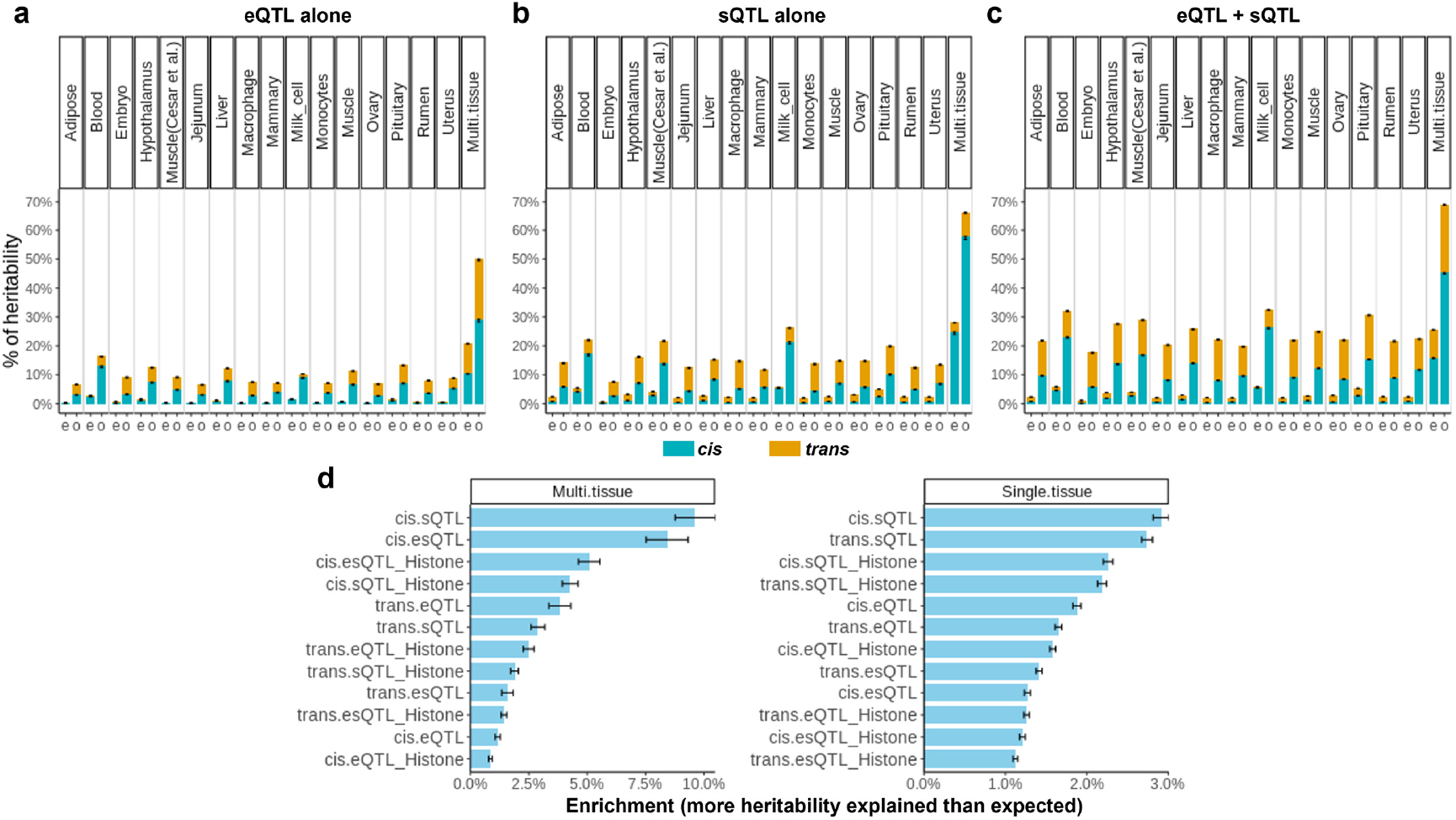
Proportions of genetic variance or heritability explained by regulatory variants. **a**: when only eQTL variants were considered. **b**: when only sQTL were considered. **c**: eQTL and sQTL were considered jointly. Where ‘e’ is the expected proportion of heritability explained by the genomic size, ‘o’ is the observed proportion of heritability and Multi.tissue is the regulatory variants detected from 16 tissues. Means and standard error bars across 37 traits are presented. **d**: Enrichment of heritability across fitted classes in the joint model (12 regulatory classes). The enrichment was calculated as the difference between the observed proportion of heritability explained and the expected proportion of heritability explained from the number of variants in each class. The label ‘_Histone’ means that eQTL or sQTL were also under histone marks tagged by ChIP-seq peaks.

By ranking classes of variants based on the difference between the observed and expected proportion of heritability explained (Figure 1d), multi-tissue *cis* sQTL and *cis* esQTL (variants are both eQTL and sQTL) explained the most additional variance. Multi-tissue *trans* eQTL, sQTL and esQTL also explained more heritability than expected from the number of variants in each class. At the single-tissue level, *cis* and *trans* sQTL had the greatest additional variance explained. Across all scenarios, e/sQTL under histone marks tagged by ChIP-seq peaks explained more heritability than expected, but not necessarily more than e/sQTL outside of histone marks (Figure 1d).

BayesRC estimated the number of trait-associated variants, i.e., QTL, with small-, medium- and large-effect within each regulatory class. We then compared the proportion of variants within each class that fell into the small, medium and large-effect QTL distributions. By comparing each regulatory class to the remaining class (no regulatory evidence) we estimated the additional proportion of QTL in each class than expected by the number of variants of that class (Figure 2a). Across analysed traits, multi-tissue *cis* eQTL had the most additional proportion of QTL than expected. Driven by the relatively small number of variants identified (e.g., Table 1), *trans* e/sQTL also had a high additional proportion of QTL than expected in both single-tissue and multi-tissue analyses. Regulatory variants that were either eQTL or sQTL and under histone marks had the greatest additional proportion of QTL than expected, suggesting combining regulatory classes improves the mapping of potentially causal variants.

**Figure 2.**
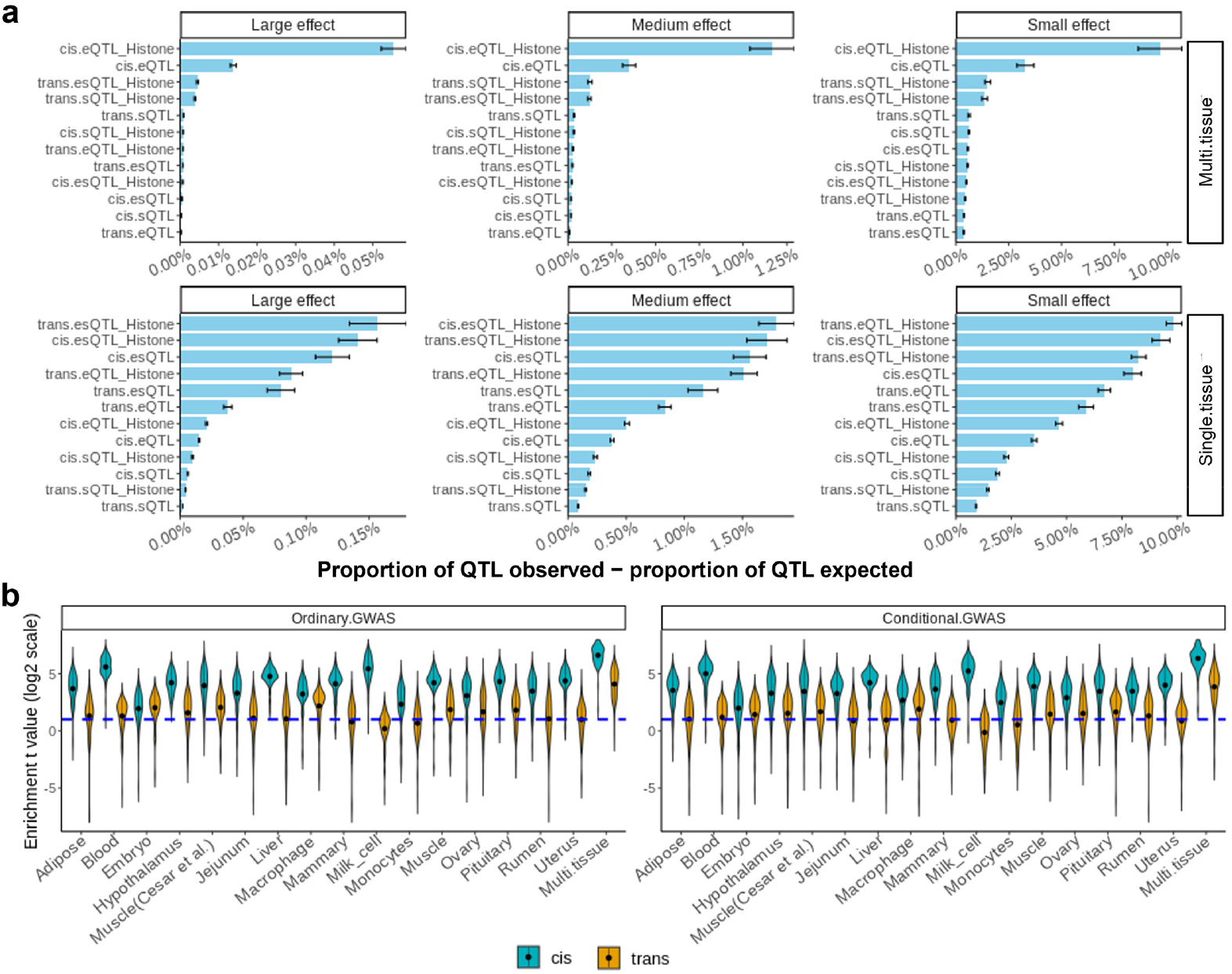
Amount of trait-associated variants (QTL) included in regulatory variants. **a**: For each class, the difference between the observed proportion (or concentration) of QTL and the expected proportion of QTL by genome size in the BayesRC analysis is the additional proportion of QTL included. The label ‘_Histone’ means that eQTL or sQTL were also under histone marks tagged by ChIP-seq peaks. **b**: The enrichment of QTL in regulatory variants was determined as the difference (t value) of variant effects in GWAS between a set of e/sQTL and a set of random variants with matched LD and MAF to the e/sQTL set. The blue dashed line indicates t value = log(2) which is equivalent to the p-value threshold of 0.05. Each violin bar represents the results across 37 traits. “Conditional.GWAS”: GWAS results conditioned on the top-2 QTL per chromosome from the results of the ordinary GWAS (“Ordinary.GWAS”, no top QTL was fitted).

It is possible that the classes of regulatory variants differ in minor allele frequency (MAF) or LD and that this explains the enrichment of QTL and genetic variance within regulatory classes. To test this possibility, we implemented a MAF-LD matched enrichment test (see Methods) using GWAS results of 37 traits on 16 million sequence variants ^8^. For each class of regulatory variants, e.g., cis eQTL, we sampled a random set of variants (repeated 1000 times) with matched MAF and LD, then we compared the GWAS effects between the set of regulatory variants and the set of random variants with matched MAF and LD. To ensure that the results are not driven by a few large-effect QTL, we carried out another set of GWAS of the 37 traits conditional on the effects of the top 2 variants per chromosome at least 1Mb apart (i.e. we fitted the top 2 variants per chromosome in the statistical model as fixed effects, see Methods). We then applied the MAF-LD matched enrichment test to the conditional GWAS. As shown in Figure 2b, across traits and tissues, both proximal and distal regulatory variants were significantly enriched with QTL compared to random variants with matched MAF and LD using both the original or conditional GWAS. The strongest enrichment of QTL was found in e/sQTL from multiple tissues. Therefore, these results confirmed that the enrichment of QTL in regulatory variants was not driven by MAF or LD.

We next examined whether the contribution of regulatory variants to trait heritability was consistent between different populations and could be reproduced using external datasets. As there are 37 phenotypic records on both 110k cows and 9k bulls, we conducted BayesRC fitting the 13 classes of regulatory variants separately in bulls and cows to check the variability of the enrichment of heritability between different cattle datasets. We found that the Pearson correlation of partitioned heritability across 12 regulatory classes from different tissues for 37 traits was 0.85, with the correlation for results from single and multiple tissues being 0.846 and 0.824, respectively (Figure 3a).

**Figure 3.**
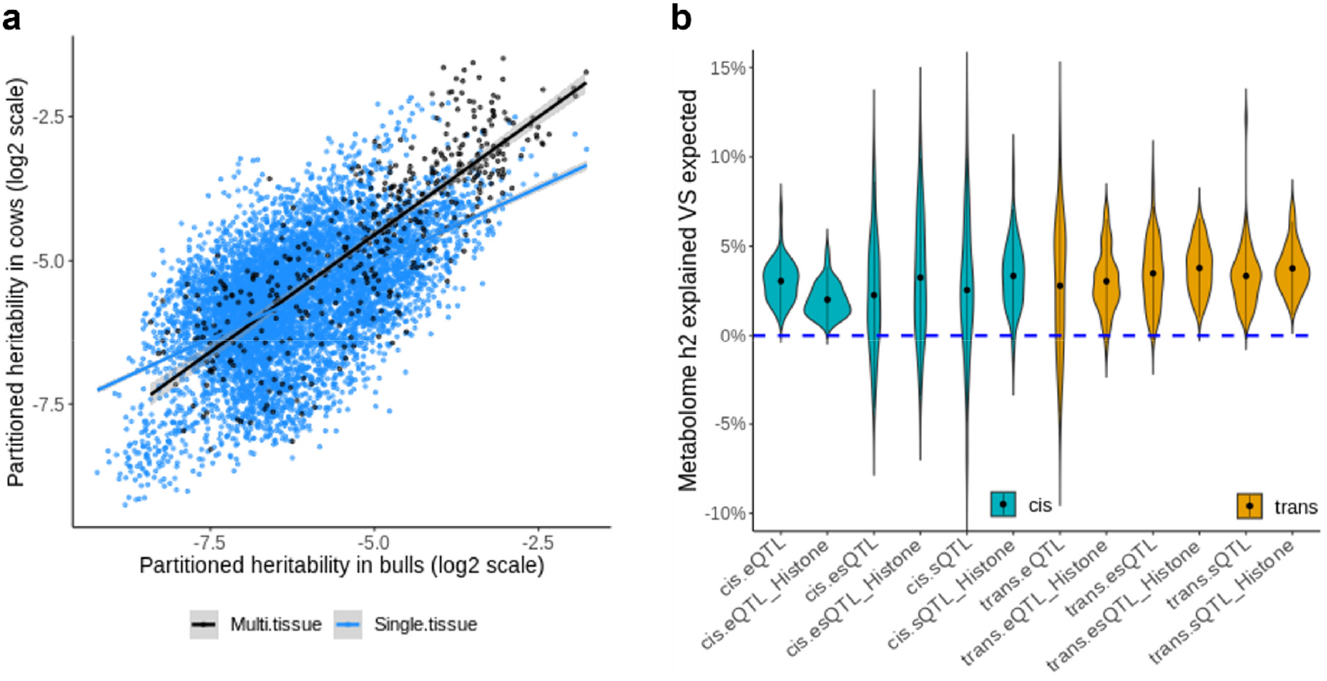
Consistency and metabolic analysis of heritability explained by regulatory variants. **a**: A scatter plot of the heritability explained by 12 classes of regulatory variants across tissues and traits between bulls and cows. The black and blue lines are the regressions for heritability partitioned using multi-tissue and single-tissue regulatory variants, respectively. **b**: The proportion of heritability explained by multi-tissue regulatory variants more than expected by genomic size averaged across 56 metabolic traits and different classes.

To further verify the large proportion of heritability explained by regulatory variants, we used multi-tissue e/sQTL to define classes for BayesRC to partition heritability in the metabolome, including 56 metabolomic (polar lipid) traits assayed by liquid chromatography-mass spectrometry (LCMS) on 320 cattle (see Methods). Across these 56 metabolic traits, on average *cis* and *trans* e/sQTL together explained 71.5% (SE=0.3%) of the heritability, 36.6% (0.6%) more than expected if the regulatory variants explained as much genetic variance per variant as the variants that are neither eQTL nor sQTL. Both *cis* and *trans* e/sQTL contribute substantially to the heritability of metabolic profiles (Figure 3b). A full list of partitioned heritability for the metabolic phenotypes can be found in Supplementary Data 2.

In Figure 4 we provide examples where *cis* or *trans*-regulatory variants significantly affect complex traits and are also supported by external functional information. If a causal variant is in very high LD with 3 other non-causal variants (not unlikely among 1.8 M variants), then the BayesRC analysis is likely to estimate the posterior inclusion probability (PIP) of all 4 variants to be 0.25. Therefore, we considered variants with PIP>0.25 as potentially causal. For instance, we highlight a *cis* eQTL from blood at Chr15:42044576 (rs137255300) which affected both the birth size and the concentration of lactosylceramide in cattle (Figure 4a left and middle panels). Chr15:42044576 is a missense mutation^17^ for *IRAG1* and conserved across 100 vertebrates (PhastCon score = 0.999), but this mutation affects the expression of *CTR9* (right panel), which is a transcription factor. Another example is a multi-tissue *trans* eQTL (Chr5:105773809, rs109676906) which significantly affects cattle height (Figure 4b). This single mutation explained ~0.6% of the phenotypic variance of stature in an additional 133,306 cattle across more than 19 populations/breeds ^7,18^. A list of *cis* and *trans* e/sQTL affecting different complex and metabolic traits with their functional annotation is provided in Supplementary Data 3.

**Figure 4.**
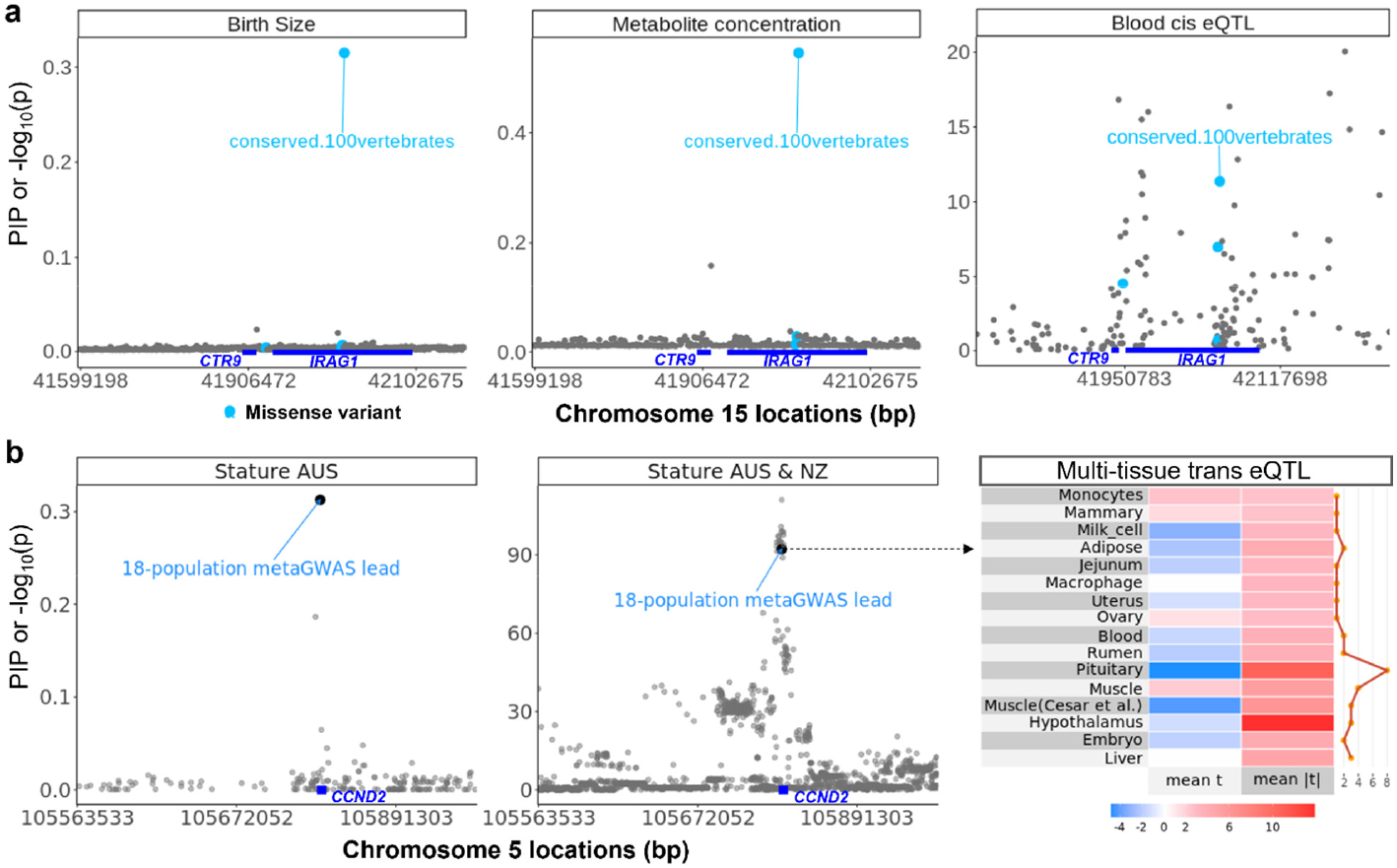
Examples of *cis* and *trans* eQTL affecting complex traits. **a**: A candidate causal mutation (Chr15:42044576, rs137255300) within *IRAG1* for birth size (left panel) and the concentration of lactosylceramide (middle panel) is a *cis* eQTL for *CTR9* in blood (right panel). Chr15:42044576 is also a missense mutation for *IRAG1* at a site conserved across 100 vertebrates. The y-axis of the left and middle panels are posterior inclusion probability (PIP) of BayesRC and the -log10(p) of eQTL mapping in blood in the right panel. **b**: A candidate causal mutation (Chr5:105773809, rs109676906) within *Cyclin D2* (*CCND2*) for stature (black point in left and middle panels) is a *trans* eQTL across multiple genes and tissues (right panel). Chr5:105773809 is also a lead variant in a meta GWAS of cattle stature across 18 global populations ^18^. The y-axis of the left panel is the PIP of BayesRC and the y-axis of the middle panel is the -log10(p) of meta-analysis GWAS of 120k Australian and New Zealand cattle. The right panel is the heatmap of effects of the trans eQTL on the expression of genes averaged within each tissue, where ‘mean t’ is the average t value across genes for each tissue and ‘mean Itl’ is the geometric mean of the magnitude of t values across genes for each tissue.

## Discussion

Our analysis of large datasets in cattle demonstrates that both cis and trans-regulatory variants significantly contribute to variation in complex traits. Such contribution is not due to the LD or MAF of regulatory variants and increases when more regulatory variants of different types (e.g., eQTL and sQTL) and a large number of tissues are included in the analysis. When *cis* and *trans* eQTL and sQTL from multiple tissues are jointly analysed, they accumulatively explain the majority (~70%) of heritability. Therefore, we expect that as more regulatory variants are discovered from more assays, tissues and individuals they will explain an even larger proportion of the heritability of complex traits.

Our study highlights the importance of sample size in the e/sQTL mapping and the detection of their overlaps with QTL. Compared to the previous cattle study ^12^ where cis eQTL contributed around 10% of heritability, the current study had increased the sample sizes for the mapping of e/sQTL by up to 20-fold (N~205 VS N~4725). The current study also increased the sample size of the mapping of complex traits by 2.5-fold which increases the power of BayesRC analysis.

Our findings contrast with several recent studies on humans where regulatory variants such as eQTL contribute a small part to phenotypic variation ^9,10,19^. Our analysis supporting the direct role of regulatory variants in shaping complex traits has several differences from previous studies which may have led to our different conclusions. One obvious distinction is that cattle are a different species from humans, although previous studies showed high similarities in genomic features between these two species ^18,20^. It is also worth noting that there are usually many more disease-related traits that underwent purifying selection in humans than in cattle.

The second distinction of our study is that when analysing variant-trait associations, we used Bayesian methods. Our BayesRC^13^ analysis used raw data that fits all variants simultaneously while most human studies use GWAS or summary statistics of GWAS (e.g., Yao et al ^9^) which associates one variant at a time with the phenotype. BayesRC ^13^ selects the variants to include in the model and estimates their effects jointly. It also allows the distribution of effects to vary between classes and fits the different class annotations jointly in the model. When similar Bayesian methods were used in human datasets ^21,22^ they showed better performances in training genomic predictors than using GWAS results. However, these Bayesian analyses did not fit different distributions of variant effect to different classes of regulatory variants. In addition, raw data is more powerful than summary statistics, where they are available.

The third distinction is that we jointly modelled multiple categories of regulatory variants, including eQTL and sQTL and variants under histone marks from multiple tissues. Although sQTL were first discovered to be important to complex traits in humans ^3^, they have not always been analysed together with eQTL in human studies of the phenotypic effects of regulatory variants ^9,10,23,24^. The current study observed that at the same p-value threshold, a lot more sQTL (3 times more in single-tissue analysis) were called than eQTL and therefore, they alone or in combination with eQTL explained more heritability than eQTL alone. In fact, multi-tissue sQTL alone explained a similarly large proportion (66%) of heritability to the proportion of heritability explained jointly by eQTL and sQTL (70%). This again validates the important role of sQTL in shaping mammalian complex traits.

The fourth difference between this study and most others is that we included *trans* eQTL and sQTL whereas most only included *cis*. In the human GTEx analysis ^25^ only a few *trans* eQTL were identified and this may have limited their use in the downstream analysis. Due to the small effect size, *trans* eQTL mapping requires a large sample size but the accumulated phenotypic effects of them may be more estimable. The CattleGTEx had different individuals per tissue which means the total sample size approaches 5000 in the multi-tissue analysis. Discovered from the CattleGTEx population and tested in the Australian population, on average single-tissue *trans* e/sQTL explained 12% of heritability and multi-tissue *trans* e/s QTL explained 24% of heritability. These findings demonstrate the important role of distal regulatory variants in shaping complex traits.

To further validate the contribution of regulatory variants to phenotypes we applied the same BayesRC methods fitting the multi-tissue e/sQTL data as biological priors to a set of metabolic phenotypes. These traits, where large effect QTL exist, are genetically simpler than traits like milk production or body size ^12,26^. We found that more than 70% of heritability in the metabolic phenotypes could be explained by both *cis* and *trans*-regulatory variants. One highlighted example is *cis* eQTL Chr15:42044576 (rs137255300) which affected both the birth size and the concentration of lactosylceramide (Figure 5a). It’s causal candidacy for these two traits is supported by external functional annotation as it is also a missense mutation and at conserved sites across 100 vertebrates. It is worth noting that Chr15:42044576 is a missense mutation for *IRAG1* but affected the expression of the nearby transcription factor gene *CTR9,* which appears to show bystander effects like *FTO* ^27,28^. This implies complex consequences of large-effect mutations on both activities of protein-coding and transcription. We also highlight a multi-tissue *trans* eQTL Chr5:105773809 (rs109676906) within *CCND2* affecting cattle stature. This mutation is not at a conserved site but had a large and replicable effect on stature in ~200,000 cattle across 19 populations across the globe ^18^. Its effect on gene expression in different tissues tended to have different directions (Figure 5b) which is consistent with the expectation of effect patterns of *trans* eQTL ^1^.

Taken together, using cattle as a model, we demonstrate the significant and direct role of *cis* and *trans*-regulatory variants in shaping mammalian complex traits. Our findings suggest that many QTL have an impact on the regulation of transcription. Therefore, with proper analysis and sufficient power, regulatory variants not only provide etiology behind the genome-to-phenome map but also are a powerful resource to directly map causal variants for mammalian complex traits.

## Methods

### RNA-seq data

The RNA-seq and genotype data analysed included those generated by Agriculture Victoria Research (AVR) in Victoria, Australia, and those provided by the cattle CattleGTEx consortium ^4^ (Supplementary Table 1). The animal ethics was approved by the DJPR Animal Ethics Committee (application numbers 2013-14 and 2018-2019), Australia. Blood samples were taken from 390 lactating cows from 2 breeds, and milk samples from 281 lactating cows from 2 breeds. The processing of samples, RNA extractions, and library preparation followed that previously described ^29,30^. RNA sequencing (RNA-seq) was performed on a HiSeq3000 (Illumina Inc) or NovaSeq6000 (Illumina Inc) genome analyzer in a paired-end, 150-cycle run. Only RNA-seq data of 356 Holstein and 26 Jersey with > 50 million reads for milk cells or > 25 million reads for white blood cells and had concordant alignment rate ^31^ > 80% were used. QualityTrim (https://bitbucket.org/arobinson/qualitytrim) was used to trim and filter poor-quality bases and sequence reads. Adaptor sequences and bases with a quality score of <20 were removed. Reads with a mean quality score less than 20, greater than 3 N, greater than three consecutive bases with a quality score less than 15, or a final length of fewer than 50 bases were discarded. High-quality raw reads were aligned to the ARS-UCD1.2 bovine genome ^32^ with STAR ^31^ using the 2-pass method. The gene counts were extracted by FeatureCount ^33^. Leafcutter ^34^ was used to generate junction files which were then used to create the RNA splicing phenotype matrix, i.e., intron excision ratio ^34^.

The RNA-seq gene counts of 15 tissues (Supplementary Table 1) where the sample size > 100 were downloaded from CattleGTEx website http://cgtex.roslin.ed.ac.uk/. The blood counts generated by AVR and CattleGTEx were combined. All gene counts were normalised by voom ^35^ and then underwent quantile normalisation for the following analyses. Junction files from CattleGTEx tissues were also downloaded and data from each tissue were processed by leafcutter ^34^ to generate RNA splicing phenotype. Milk cell data used in this study was only from AVR.

### Genotype data

The genotype data for Australian animals including those used for e/sQTL mapping (blood and milk cells) and association analysis of phenotypes (described later) were 16,251,453 sequence variants imputed using Run7 of the 1000 Bull Genomes Project ^36,37^. The details of the imputation were described previously ^38^. Briefly, the imputation of biallelic sequence variants was performed with Minimac3 ^39,40^ and those variants with imputation accuracy *R^2^* > 0.4 and minor allele frequency (MAF) > 0.005 in both bulls and cows were kept. Bulls were genotyped with either a medium-density SNP array (50K: BovineSNP50 Beadchip, Illumina Inc) or a high-density SNP array (HD: BovineHD BeadChip, Illumina Inc) and cows were genotyped with the BovineSNP50 Beadchip (Illumina Inc). The genotype data for CattleGTEx animals were generated previously ^4^ and included a total of more than 6 million sequence variants imputed also using Run7 of the 1000 Bull Genomes Project. Those variants with the imputation dosage R-squared > 0.8 and MAF > 0.001 were kept.

### Phenotype data

Data were collected by farmers and processed by DataGene Australia (http://www.datagene.com.au/) for the official May 2020 release of National breeding values. No live animal experimentation was required. DataGene provided the bull and cow phenotypes as de-regressed breeding values or trait deviations for cows, and daughter trait deviations for bulls (ie. progeny test data for bulls). DataGene corrected the phenotypes for herd, year, season and lactation following the procedures used for routine genetic evaluations in Australian dairy cattle. Phenotype data included a total of 8,949 bulls and 103,350 cows, including Holstein (6,886♂ / 87,003♀), Jersey (1562♂ / 13,353♀), cross-breed (36♂ / 5,037♀) and Australian Red (265♂ / 3,379♀) dairy breeds. In total, 37 traits were studied that related to milk production, mastitis, fertility, temperament and body conformation and the details of these traits can be found in ^38^. For AVR blood samples breed and days in milk (DIM) were fitted as fixed effects in the model. For the milk samples, experiment, DIM and the first and second principal components, extracted from the expression count matrix, were fitted as fixed effects. This aimed at adjusting the high expression of casein genes in milk cells based on previous experiences ^29^.

### Mapping and selection of eQTL and sQTL

A GWAS approach that fits random effects of a relationship matrix ^14^ can control false correlations and that was used in the current study to map eQTL and sQTL:

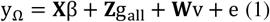

Where y_Ω_ is an n × 1 vector of omics values such as gene expression or RNA splicing, **X** was the design matrix allocating phenotypes to fixed effects; *β* is a vector of fixed effects like breeds, different experiments, or PEER ^41^ factors derived by the CattleGTEx consortium ^4^, ***Z*** is a matrix allocating records to individuals; *g*_Ω_*all*__ is an n × 1 vector of the total genetic effects of the individuals with 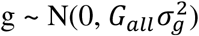 where *G_all_* is the genomic relationship matrix (GRM) built by all the variants; **W** is the design matrix of variant genotypes (0, 1, 2) and *ν* is the variant additive effect; e is the error term.

*cis* e/sQTL were defined as those variants within ±1Mb of the transcription start site of a gene or down/upstream of an intron with p < 5e-6 in GWAS. This threshold resulted in that, on average across tissues, the false discovery rate (FDR) was 0.0158(6e-5) for eQTL mapping and 0.0164(2e-5) for sQTL mapping (see equation 2 described in the following). Trans e/sQTL were defined as those not on the same chromosome of the omics feature with p < 5e-6 in GWAS. Only the top 3 trans e/sQTL per chromosome were selected. In addition, we impose (FDR) of:

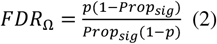

where *p* is the GWAS p-value cutoff, e.g., 5e-6, *Prop_sig_* is the proportion of variants significant given the GWAS p-value cutoff to the total number of variants analysed. If *FDR*_Ω_ >= 0.05 for an omics feature, no trans e/sQTL were selected. Also, those e/sQTL under at least two ChIP-seq peaks identified from multiple studies ^6,42,43^ were used in the Bayesian analysis described below.

### Meta-analysis of e/sQTL

Because data from different tissues of CattleGTEx were from different individuals, combining results from each tissue can increase the chance of detecting causal regulatory variants. The human GTEx ^1,2^, showed that *cis* e/sQTL to a large extent showed consistent effects across tissues. Although, the ranking of the effects of the same variant across tissues may be different. For *trans* e/sQTL, it is not expected that their effects will be consistent across tissues. Considering these factors, we implemented the following 2 formulae in meta-analyses of e/sQTL:

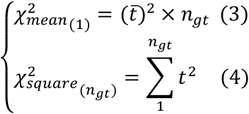

In equation 3, it is assumed that the effects of a variant across genes and tissues were largely consistent; the chi-square is based on the mean of the t value (beta/se) of variants; 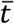 is the mean of the t-value of a variant across all genes that it affected across all tissues the effects were measured; *n_gt_* is the number of genes and tissues where the effect of this variant was estimated; 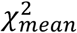 was tested against a chi-square distribution with 1 degree of freedom. In equation 4, it is not assumed that the effects of a variant across genes and tissues were largely consistent; the chi-square is based on the sum of the square of t values of variants across all genes and tissues; 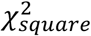 was tested against a chi-square distribution with *n_gt_* degree of freedom. For *cis* e/sQTL, both 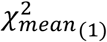 and 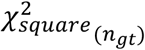 were calculated and variants with a p < 5e-8 for either 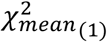 or 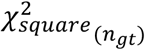 were called significant. For *trans* e/sQTL, variants with p< 5e-8 for 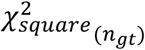 and effects estimated in at least two tissues were called significant.

### BayesRC using *cis* and *trans* e/sQTL

BayesRC ^13^ extends the classic BayesR algorithm ^15,16^ to incorporate independent classes of variants (‘*c*’) to model informative biological priors. Similar to the classic BayesR, BayesRC models the prior of variant effects which is a mixture distribution of four normal distributions including a null distribution, zero-effect [*N*(0,0.0*σ*^2^_*g*_)], and three others: small-effect [*N*(0,0.0001*σ*^2^_*g*_)], medium-effect [*N*(0,0.001*σ*^2^_*g*_)] and large-effect [*N*(0,0.01*σ*^2^_*g*_)], where *σ*^2^_*g*_ is the additive genetic variance for the trait. The BayesRC ^13^ model used here for association analysis of phenotypes was:

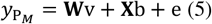

where *y*_P_*M*__ was a vector of traits, **W** was the design matrix of marker genotypes; centered and standardised to have a unit variance; *ν* was the vector of variant effects; X was the design matrix allocating phenotypes to fixed effects; *b* was the vector of fixed effects of breeds. BayesRC was conducted for 37 traits on cows and bulls separately. As a result of 50,000 iterations with 25,000 burn-ins of Markov chain Monte Carlo (MCMC), the effect ν for each variant jointly estimated with other variants was obtained. This mixture of distributions is modelled independently in each class of variants to allow for different mixture models per class (‘*c*’).

To better understand the contribution of regulatory variants to complex traits, we used different classifications to jointly or separately model eQTL and/or sQTL. When eQTL and sQTL were modelled jointly, 13 classes of variants were created (Supplementary Table 2) with the 13^th^ class being the remaining variants neither eQTL nor sQTL. When eQTL and sQTL were modelled separately, 5 classes of variants were created for eQTL and sQTL separately and the 5^th^ class was the remaining variants neither eQTL nor sQTL. Such classification, i.e., one 13-category classification and two 5-category classifications (eQTL and sQTL separately) was created for each tissue and the results of multi-tissue e/sQTL mapping. When creating these classes, variants detected as both *cis* e/sQTL and *trans* e/sQTL were set to *cis* e/sQTL. For better computational efficiency, we LD pruned (*r*^2^ < 0.9) those 16 million variants using plink ^44^ and used the resultant 1,882,504 variants for BayesRC.

### Partitioning heritability across functional classes

MCMC in BayesRC estimated additive genetic variance (Va) based on sequence variants and the total error variance (Ve) and this can be used to calculate the heritability of each trait 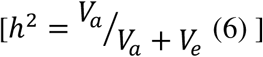. Results from BayesRC from cows and bulls were both analysed and the average between the two estimates was presented. MCMC in BayesRC also estimated the number of variants in each class (e.g., *cis* eQTL, *trans* eQTL) that fell into the 4 distributions of effects: zero-effect [*N*(0,0.0*σ*^2^_*g*_) |, small-effect [*N*(0,0.000l*σ*^2^_*g*_)], medium-effect pV(0,0.001 *σ*^2^_*g*_)] and large-effect [*N*(0,0.0l*σ*^2^_*g*_)], where *σ*^2^_*g*_ was the additive genetic variance for the trait. This can be used to partition Va and thus, *h*^2^, into each class:

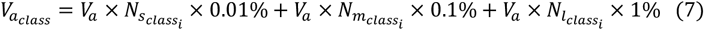

Where *N_s_class_i___* was the number of small-effect variants in class *i* (e.g., *cis* eQTL), *N_m_class_i___* was the number of medium-effect variants in *cis* e/sQTL and *N_l_class_i___*. was the number of large-effect variants in *cis* e/sQTL. Then for each class, we used equation 6 to calculate *h*^2^ for each class 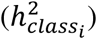, and then the proportion of *h*^2^ explained by each class as: 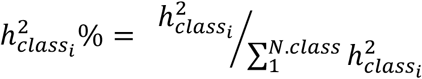 (equation 8) where *N. class* was the total number of classes fitted in the model.

We derive an expected 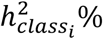, or 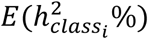 using the 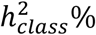 and the proportion of variants for the remaining class (variants were neither eQTL nor sQTL):

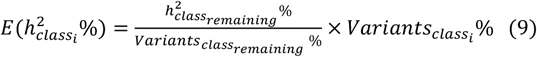

where 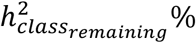 was the proportion of heritability explained by the class of remaining variants, *Variants_class_remaining__* % was the proportion of the class of remaining variants to the total number of variants analysed and *Varaiants_class_i__*% was the proportion of the class *i* of variants (e.g., *cis* eQTL) to the total number of variants analysed. When 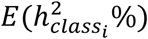was derived, 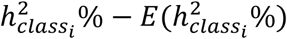 can be used to estimate the amount of heritability explained by each class as a deviation from that expected by the size of class *i*.

Applying the same mechanism as above, we estimated the expected proportion of trait-associated variants (QTL) for each class:

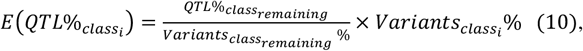

where *QTL%_class_remaining__* was the proportion of QTL in the class of remaining variants. Then *QTL%_class_i__*. – *E*(*QTL*%_*class*_*i*__) can be used to estimate the proportion of QTL included in each class as a deviation from that expected by the size of class *i*.

### MAF-LD matched enrichment test

Using the Australian cattle genotype data the 16 million sequence variants were first divided into 20 bins using LD score (50kb window size) calculated using GCTA ^14^. Within each of these LD bins, we then divided variants into 20 bins of MAF. This divided the 16 million variants into 400 LD-MAF bins. Then, for a given set of regulatory variants, e.g., *cis* eQTL from blood, we laid them over 400 LD-MAF bins to identify LD-MAF bins associated with this set of regulatory variants and the number of regulatory variants falling into each bin (*N_reg_bin_i___*). Within each of these LD-MAF bins associated with the regulatory variants, we sampled a random set of *N_reg_bin_i___* variants. This random sampling was repeated 1000 times. For the set of regulatory variants, we used the significance from the GWAS and conditional GWAS (detailed in next paragraph), i.e., -log(GWAS p), to indicate the effect size which was averaged across all regulatory variants. Then, for each of 1000 sets of LD-MAF matched random variants, the average -log(GWAS p) was also calculated. We then used a t-test to quantify the difference of 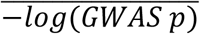 between regulatory variants and LD-MAF matched random variants, where we used the t value to indicate the enrichment of GWAS hits in regulatory variatnts compared to that expected by random variants with matched LD and MAF.

### GWAS and conditional GWAS were used for the enrichment test

The original GWAS of 37 traits in cows had been conducted previously ^8^. Briefly, the following linear mixed model was used:

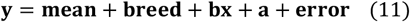

where **y** = vector of phenotypes for bulls or cows, breed = four breeds for cows (Holstein, Jersey, Australian Red and MIX); **bx** = regression coefficient *b* on variant genotypes **x**; **a** = random polygenic effects 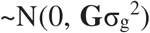 where **G** = genomic relatedness matrix based on all variants and 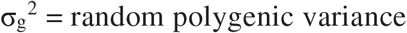; **error** = the vector of random residual effects 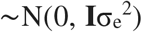, where I = the identity matrix and 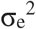 the residual variance. The construction of GRM followed the default setting (--make-grm) in GCTA^14^.

The above described MAF-LD matched enrichment test used both the original and conditional GWAS. The purpose of using conditional GWAS to conduct the enrichment test was to make sure that the enrichment was not driven by a few large-effect QTL on each chromosome. We first selected the top 2 variants based on the p-value of the original GWAS on each chromosome which were at least 1Mb apart. Then we fit these ~2 × 30 top variants in the COJO analysis implemented in GCTA ^45^ to obtain GWAS results conditioned on these top variants for 37 traits. Then the MAF-LD matched enrichment test was applied to the results of conditional GWAS of 37 traits.

### Metabolomics QTL

The discovery of milk fat polar lipid metabolite QTLs (mQTLs) was based on the mass-spectrometry quantified concentration of 59 polar lipids in milk from 338 Holstein cows (Supplementary Table 5). The bovine milk was collected as described previously ^29^ and polar lipids were extracted from bovine milk following the previously developed protocols ^46^. The chromatographic separation of polar lipids used a Luna HILIC column (250×4.6 mm, 5 μm, Phenomenex) maintained at 30 °C. The lipids were detected by the LTQ-Orbitrap mass spectrometer (Thermo Scientific) operated in electrospray ionization positive (for most polar lipid classes) or negative (for analysis of PI) Fourier transform mode. The identification of lipid species present in milk was performed as previously reported ^46^. Quantification of selected polar lipid species was based on the peak area of parent ions after normalization by the internal standard. After a quality check, data from 56 metabolites from 320 cows were used for further analysis.

We applied the same BayesRC model in equation 5 to analyse each of these metabolites, with additional fixed effects of year and batch. The biological prior for the analysis of metabolites used the 13 classes of regulatory variants detected from multiple tissues as this set explained the largest proportion of heritability for conventional traits. Then we applied equations 6-8 to partition the heritability of metabolic traits. We raised the MAF cutoff to >0.025 in the analysis of metabolic traits as the sample size is relatively small.

### Conserved variants

Conserved genome sites in cattle were based on the lifted over (https://genome.ucsc.edu/cgi-bin/hgLiftOver) human sites with PhastCon score ^47^ >0.8 computed across 30 mammals and 100 vertebrate species. The human PhastCon data was downloaded from UCSC genome database (http://hgdownload.cse.ucsc.edu/goldenpath/hg38/phastCons30way/ and http://hgdownload.cse.ucsc.edu/goldenpath/hg38/phastCons100way/). The downloaded Wiggle files were converted to bed files which were used by the LiftOver tool as an input. Another input for LiftOver was the chain file between hg38 and cattle ARS-UCD1.2.

### Meta-analysis of GWAS

For variants that appeared in multiple studies, we used the formula based on the inversed variance from METAL ^48^ to conduct meta-analysis. When combined betameta and semeta were obtained we calculated the t_meta_ = beta_meta_ / se_meta_ and the phenotypic variance explained by a variant was determined by the formula:

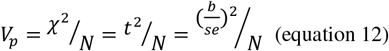

where where *V_p_* was the proportion of phenotypic variance explained by a variant, *χ^2^* was the chi-square value of the effect of the variant which is equal to the square of t value (b/se), *t*^2^, of the effect of the variant from GWAS; N was the sample size of the GWAS; b was the GWAS beta of the variant and se is the standard error of b.

## Data and code availability

The newly generated RNA-seq data (356 blood and 268 milk cells) will be made public via NCBI SRA (accession available upon manuscript publication). Other RNA-seq data can be accessed via the CattleGTEx consortium (http://cgtex.roslin.ed.ac.uk/). Linear mixed modelbased summary statistics of mapped eQTL and sQTL from each of the 16 tissue and the multi-tissue analysis is available at https://figshare.com/s/d6582f654a8a25160946. The DNA sequence data as part of the 1000 Bull Genomes Consortium^49,50^ are available to consortium members and the membership is open. Sequence data of 1832 samples from the 1000 Bull Genome Project have been made publicly available at https://www.ebi.ac.uk/eva/?eva-study=PRJEB42783. DataGene Australia (http://www.datagene.com.au/) are custodians of the raw phenotype and genotype data of Australian farm animals. Access to these data for research requires permission from DataGene under a Data Use Agreement. Other supporting data are shown in the Supplementary Materials of the manuscript. The linear mixed model analysis used GCTA ^14^. The Bayesian analysis used BayesRC^13^.

## Acknowledgments

Australian Research Council’s Discovery Projects (DP160101056 and DP200100499) supported R.X., M.E.G. and N.R.W. DairyBio, a joint venture project between Agriculture Victoria (Melbourne, Australia), Dairy Australia (Melbourne, Australia) and the Gardiner Foundation (Melbourne, Australia), funded computing resources used in the analysis. N.R.W. acknowledged funding from National Health and Medical Research Council (NHMRC 1113400 and 1078901). L.F. received funding from European Union’s Horizon 2020 research and innovation programme under the Marie Skłodowska–Curie grant (agreement No. 801215). A. T. acknowledged funding from the BBSRC through programme grants BBS/E/D/10002070 and BBS/E/D/30002275, MRC research grant MR/P015514/1, and HDR-UK award HDR-9004. The authors also thank the University of Melbourne, Australia for supporting this research. No funding bodies participated in the design of the study nor analysis, interpretation of data, or writing the manuscript. The 1000 Bull Genomes consortium provided access to cattle sequence data and the CattleGTEx consortium provided access to expression data.

## Author contributions

R.X. and M.E.G. conceived the study. R.X. carried out the main analyses with assistance from M.E.G. and E.J.B.. L.F., S.L., Y.G., G.E.L and A.T. assisted in the analysis of data from CattleGTEx. Z.L. generated the metabolomics data and I.M.M. assisted in the analysis of the metabolomics data. B.A.M. and A.J.C. generated and assisted in the analysis of the new RNA-seq data. M.E.G. and N.R.W. oversaw the project. R.X. and M.E.G. wrote the paper. R.X., M.E.G., N.R.W., L.F., S.L., A.T., A.J.C. revised the paper. All authors read and approved the final manuscript.

## Notes

### Competing Interest Statement

The authors have declared no competing interest.

## References

1 Consortium, G. The Genotype-Tissue Expression (GTEx) pilot analysis: multitissue gene regulation in humans. Science 348, 648–660 (2015).

2 Consortium, G. The GTEx Consortium atlas of genetic regulatory effects across human tissues. Science 369, 1318–1330 (2020).

3 Li, Y. I. et al. RNA splicing is a primary link between genetic variation and disease. Science 352, 600–604, doi:10.1126/science.aad9417 (2016).

4 Liu, S. et al. A comprehensive catalogue of regulatory variants in the cattle transcriptome. bioRxiv, 2020.2012.2001.406280, doi:10.1101/2020.12.01.406280 (2021).

5 Clark, E. L. et al. From FAANG to fork: application of highly annotated genomes to improve farmed animal production. Genome Biology 21, 1–9 (2020).

6 Kern, C. et al. Functional annotations of three domestic animal genomes provide vital resources for comparative and agricultural research. Nature Communications 12, 1821, doi:10.1038/s41467-021-22100-8 (2021).

7 Reynolds, E. G. et al. Non-additive association analysis using proxy phenotypes identifies novel cattle syndromes. Nature Genetics, 1–6 (2021).

8 Xiang, R. et al. Mutant alleles differentially shape fitness and other complex traits in cattle. Communications Biology 4, 1–10 (2021).

9 Yao, D. W., O’Connor, L. J., Price, A. L. & Gusev, A. Quantifying genetic effects on disease mediated by assayed gene expression levels. Nature Genetics 52, 626–633 (2020).

10 Connally, N. et al. The missing link between genetic association and regulatory function. medRxiv (2021).

11 van den Berg, I. et al. Meta-analysis for milk fat and protein percentage using imputed sequence variant genotypes in 94,321 cattle from eight cattle breeds. Genetics Selection Evolution 52, 37, doi:10.1186/s12711-020-00556-4 (2020).

12 Xiang, R. et al. Quantifying the contribution of sequence variants with regulatory and evolutionary significance to 34 bovine complex traits. Proceedings of the National Academy of Sciences 116, 19398–19408 (2019).

13 MacLeod, I. et al. Exploiting biological priors and sequence variants enhances QTL discovery and genomic prediction of complex traits. BMC genomics 17, 144 (2016).

14 Yang, J., Lee, S. H., Goddard, M. E. & Visscher, P. M. GCTA: a tool for genome-wide complex trait analysis. The American Journal of Human Genetics 88, 76–82 (2011).

15 Erbe, M. et al. Improving accuracy of genomic predictions within and between dairy cattle breeds with imputed high-density single nucleotide polymorphism panels. Journal of dairy science 95, 4114–4129 (2012).

16 Moser, G. et al. Simultaneous discovery, estimation and prediction analysis of complex traits using a Bayesian mixture model. PLoS genetics 11, e1004969 (2015).

17 McLaren, W. et al. The Ensembl Variant Effect Predictor. Genome Biology 17, 122, doi:10.1186/s13059-016-0974-4 (2016).

18 Bouwman, A. C. et al. Meta-analysis of genome-wide association studies for cattle stature identifies common genes that regulate body size in mammals. Nature genetics 50, 362 (2018).

19 Mostafavi, H., Spence, J. P., Naqvi, S. & Pritchard, J. K. Limited overlap of eQTLs and GWAS hits due to systematic differences in discovery. bioRxiv (2022).

20 Liu, S. et al. Epigenomics and genotype-phenotype association analyses reveal conserved genetic architecture of complex traits in cattle and human. BMC biology 18, 1–16 (2020).

21 Lloyd-Jones, L. R. et al. Improved polygenic prediction by Bayesian multiple regression on summary statistics. bioRxiv, 522961 (2019).

22 Patxot, M. et al. Probabilistic inference of the genetic architecture underlying functional enrichment of complex traits. Nature Communications 12, 1–16 (2021).

23 Boyle, E. A., Li, Y. I. & Pritchard, J. K. An expanded view of complex traits: from polygenic to omnigenic. Cell 169, 1177–1186 (2017).

24 Võsa, U. et al. Large-scale cis-and trans-eQTL analyses identify thousands of genetic loci and polygenic scores that regulate blood gene expression. Nature genetics 53, 1300–1310 (2021).

25 Consortium, G. Genetic effects on gene expression across human tissues. Nature 550, 204 (2017).

26 Liu, Z. et al. Fine-mapping sequence mutations with a major effect on oligosaccharide content in bovine milk. Scientific reports 9, 1–12 (2019).

27 Xiang, R., van den Berg, I., MacLeod, I. M., Daetwyler, H. D. & Goddard, M. E. Effect direction meta-analysis of GWAS identifies extreme, prevalent and shared pleiotropy in a large mammal. Communications biology 3, 1–14 (2020).

28 Claussnitzer, M. et al. FTO obesity variant circuitry and adipocyte browning in humans. New England Journal of Medicine 373, 895–907 (2015).

29 Xiang, R. et al. Genome variants associated with RNA splicing variations in bovine are extensively shared between tissues. BMC Genomics 19, 521, doi:10.1186/s12864-018-4902-8 (2018).

30 Chamberlain, A. et al. in 11th world congress on genetics applied to livestock production (WCGALP). Auckland, New Zealand: Volume Molecular Genetics. 254.

31 Dobin, A. et al. STAR: ultrafast universal RNA-seq aligner. Bioinformatics 29, 15–21, doi:10.1093/bioinformatics/bts635 (2013).

32 Rosen, B. D. et al. De novo assembly of the cattle reference genome with single-molecule sequencing. GigaScience 9, giaa021 (2020).

33 Liao, Y., Smyth, G. K. & Shi, W. featureCounts: an efficient general purpose program for assigning sequence reads to genomic features. Bioinformatics 30, 923–930 (2014).

34 Li, Y. I. et al. Annotation-free quantification of RNA splicing using LeafCutter. Nature genetics 50, 151 (2018).

35 Law, C. W., Chen, Y., Shi, W. & Smyth, G. K. Voom: precision weights unlock linear model analysis tools for RNA-seq read counts. Genome Biology 15, R29 (2014).

36 Daetwyler, H. et al. in Proc Assoc Adv Anim Breed Genet. 201–204.

37 Daetwyler, H. et al. Integration of functional genomics and phenomics into genomic prediction raises its accuracy in sheep and dairy cattle. Proceedings of the Association for the Advancement of Animal Breeding and Genetics, Armidale, NSW, Australia, 11–14 (2019).

38 Xiang, R. et al. Mutant alleles differentially shape cattle complex traits and fitness. bioRxiv, 2021.2004.2019.440546, doi:10.1101/2021.04.19.440546 (2021).

39 Fuchsberger, C., Abecasis, G. R. & Hinds, D. A. minimac2: faster genotype imputation. Bioinformatics 31, 782–784 (2014).

40 Howie, B., Fuchsberger, C., Stephens, M., Marchini, J. & Abecasis, G. R. Fast and accurate genotype imputation in genome-wide association studies through pre-phasing. Nature genetics 44, 955 (2012).

41 Stegle, O., Parts, L., Piipari, M., Winn, J. & Durbin, R. Using probabilistic estimation of expression residuals (PEER) to obtain increased power and interpretability of gene expression analyses. Nature protocols 7, 500–507 (2012).

42 Prowse-Wilkins, C. P. et al. Putative Causal Variants Are Enriched in Annotated Functional Regions From Six Bovine Tissues. Frontiers in Genetics 12, doi:10.3389/fgene.2021.664379 (2021).

43 Xiang, R., Breen, E. J., Prowse-Wilkins, C. P., Chamberlain, A. J. & Goddard, M. E. Bayesian genome-wide analysis of cattle traits using variants with functional and evolutionary significance. Animal Production Science 61, 1818–1827, doi:https://doi.org/10.1071/AN21061 (2021).

44 Chang, C. C. et al. Second-generation PLINK: rising to the challenge of larger and richer datasets. Gigascience 4, 7, doi:10.1186/s13742-015-0047-8 (2015).

45 Yang, J. et al. Conditional and joint multiple-SNP analysis of GWAS summary statistics identifies additional variants influencing complex traits. Nature genetics 44, 369 (2012).

46 Liu, Z., Moate, P., Cocks, B. & Rochfort, S. Comprehensive polar lipid identification and quantification in milk by liquid chromatography–mass spectrometry. Journal of Chromatography B 978, 95–102 (2015).

47 Siepel, A. et al. Evolutionarily conserved elements in vertebrate, insect, worm, and yeast genomes. Genome research 15, 1034–1050 (2005).

48 Willer, C. J., Li, Y. & Abecasis, G. R. METAL: fast and efficient meta-analysis of genomewide association scans. Bioinformatics 26, 2190–2191 (2010).

49 Hayes, B. J. & Daetwyler, H. D. 1000 Bull Genomes Project to Map Simple and Complex Genetic Traits in Cattle: Applications and Outcomes. Annu Rev Anim Biosci 7, 89–102, doi:10.1146/annurev-animal-020518-115024 (2019).

50 Daetwyler, H. D. et al. Whole-genome sequencing of 234 bulls facilitates mapping of monogenic and complex traits in cattle. Nature genetics 46, 858 (2014).

