## Supplementary Table 1-5 for "Gene expression and RNA splicing explain large proportions of the heritability for complex traits in cattle"

Contains Supplementary Table 1-5.

**Supplementary Table 1**. The number of independent individuals per tissue used for eQTL and sQTL mapping.

| Tissue | N of individuals | Generation |
|---|---|---|
| Blood | 945 | AVR (356) + CattleGTEx (589) |
| Muscle | 699 | CattleGTEx |
| Liver | 576 | CattleGTEx |
| Uterus | 359 | CattleGTEx |
| Macrophage | 295 | CattleGTEx |
| Embryo | 281 | CattleGTEx |
| Milk_cell | 268 | AVR |
| Rumen | 202 | CattleGTEx |
| Mammary | 175 | CattleGTEx |
| Muscle (Cesar et al.) | 171 | CattleGTEx |
| Adipose | 151 | CattleGTEx |
| Ovary | 139 | CattleGTEx |
| Pituitary | 134 | CattleGTEx |
| Monocytes | 113 | CattleGTEx |
| Hypothalamus | 112 | CattleGTEx |
| Jejunum | 105 | CattleGTEx |
| Average | 295 | |
| Total | 4725 | |

AVR: dataset generated by Agriculture Victoria, Australia.
CattleGTEx: data from the Cattle GTEx consortium (https://www.biorxiv.org/content/10.1101/2020.12.01.406280v2).

**Supplementary Table 2.** Classification of regulatory variants based on eQTL, sQTL and ChIP-seq peaks at the single-tissue and multi-tissue level based on 1.8 million genome-wide pruned variants (LD-r2 < 0.9) in Australian cattle. esQTL had 13 classes of variants fitting eQTL and sQTL together. eQTLhad 5 classes of variants only fitting eQTL. sQTLhad 5 classes of variants only fitting sQTL. Where yesChIP were regulatory variants under ChIP-seq peaks, noChIP were regulatory variants not under ChIP-seq peaks, esQTL were regulatory variants that were both detected as eQTL and sQTL and Multi.tissue were regulatory variants detected from the multi-tissue meta-analysis.

| esQTL | | | | eQTL | | | | sQTL | |
|---|---|---|---|---|---|---|---|---|---|
| **Tissue** | **class** | **type** | **nSNP** | **Tissue** | **class** | **type** | **nSNP** | **type** | **nSNP** |
| Adipose | 1 | cis.eQTL.yesChIP | 780 | Adipose | 1 | cis.eQTL.yeschip | 1729 | cis.sQTL.yeschip | 6829 |
| Adipose | 2 | cis.eQTL.noChIP | 1396 | Adipose | 2 | cis.eQTL.nochip | 2678 | cis.sQTL.nochip | 9967 |
| Adipose | 3 | cis.sQTL.yesChIP | 5841 | Adipose | 3 | trans.eQTL.yeschip | 665 | trans.sQTL.yeschip | 10325 |
| Adipose | 4 | cis.sQTL.noChIP | 8619 | Adipose | 4 | trans.eQTL.nochip | 1605 | trans.sQTL.nochip | 24221 |
| Adipose | 5 | cis.esQTL.yesChIP | 949 | Adipose | 5 | remaining | 1875818 | remaining | 1831155 |
| Adipose | 6 | cis.esQTL.noChIP | 1282 | Blood | 1 | cis.eQTL.yeschip | 22648 | cis.sQTL.yeschip | 37584 |
| Adipose | 7 | trans.eQTL.yesChIP | 319 | Blood | 2 | cis.eQTL.nochip | 32001 | cis.sQTL.nochip | 56395 |
| Adipose | 8 | trans.eQTL.noChIP | 737 | Blood | 3 | trans.eQTL.yeschip | 1669 | trans.sQTL.yeschip | 8431 |
| Adipose | 9 | trans.sQTL.yesChIP | 9973 | Blood | 4 | trans.eQTL.nochip | 4147 | trans.sQTL.nochip | 20569 |
| Adipose | 10 | trans.sQTL.noChIP | 23340 | Blood | 5 | remaining | 1822035 | remaining | 1759516 |
| Adipose | 11 | trans.esQTL.yesChIP | 346 | Embryo | 1 | cis.eQTL.yeschip | 797 | cis.sQTL.yeschip | 466 |
| Adipose | 12 | trans.esQTL.noChIP | 868 | Embryo | 2 | cis.eQTL.nochip | 1236 | cis.sQTL.nochip | 776 |
| Adipose | 13 | remaining | 1828045 | Embryo | 3 | trans.eQTL.yeschip | 4618 | trans.sQTL.yeschip | 4334 |
| Blood | 1 | cis.eQTL.yesChIP | 9513 | Embryo | 4 | trans.eQTL.nochip | 8988 | trans.sQTL.nochip | 7481 |
| Blood | 2 | cis.eQTL.noChIP | 15338 | Embryo | 5 | remaining | 1866856 | remaining | 1869438 |
| Blood | 3 | cis.sQTL.yesChIP | 24314 | Hypothalamus | 1 | cis.eQTL.yeschip | 9412 | cis.sQTL.yeschip | 9356 |
| Blood | 4 | cis.sQTL.noChIP | 39493 | Hypothalamus | 2 | cis.eQTL.nochip | 15915 | cis.sQTL.nochip | 13653 |
| Blood | 5 | cis.esQTL.yesChIP | 13135 | Hypothalamus | 3 | trans.eQTL.yeschip | 2547 | trans.sQTL.yeschip | 14053 |
| Blood | 6 | cis.esQTL.noChIP | 16663 | Hypothalamus | 4 | trans.eQTL.nochip | 6069 | trans.sQTL.nochip | 33298 |
| Blood | 7 | trans.eQTL.yesChIP | 898 | Hypothalamus | 5 | remaining | 1848555 | remaining | 1812135 |
| Blood | 8 | trans.eQTL.noChIP | 2204 | Muscle (Cesar et al.) | 1 | cis.eQTL.yeschip | 2329 | cis.sQTL.yeschip | 24946 |
| Blood | 9 | trans.sQTL.yesChIP | 7439 | Muscle (Cesar et al.) | 2 | cis.eQTL.nochip | 3050 | cis.sQTL.nochip | 41089 |
| Blood | 10 | trans.sQTL.noChIP | 18252 | Muscle (Cesar et al.) | 3 | trans.eQTL.yeschip | 1037 | trans.sQTL.yeschip | 9335 |
| Blood | 11 | trans.esQTL.yesChIP | 771 | Muscle (Cesar et al.) | 4 | trans.eQTL.nochip | 2080 | trans.sQTL.nochip | 20106 |

| | | | | | | | | | |
|---|---|---|---|---|---|---|---|---|---|
| Blood | 12 | trans.esQTL.noChIP | 1943 | Muscle (Cesar et al.) | 5 | remaining | 1874000 | remaining | 1787021 |
| Blood | 13 | remaining | 1732535 | Jejunum | 1 | cis.eQTL.yeschip | 827 | cis.sQTL.yeschip | 3650 |
| Embryo | 1 | cis.eQTL.yesChIP | 790 | Jejunum | 2 | cis.eQTL.nochip | 1218 | cis.sQTL.nochip | 5847 |
| Embryo | 2 | cis.eQTL.noChIP | 1228 | Jejunum | 3 | trans.eQTL.yeschip | 885 | trans.sQTL.yeschip | 11403 |
| Embryo | 3 | cis.sQTL.yesChIP | 405 | Jejunum | 4 | trans.eQTL.nochip | 2025 | trans.sQTL.nochip | 24606 |
| Embryo | 4 | cis.sQTL.noChIP | 690 | Jejunum | 5 | remaining | 1877544 | remaining | 1836992 |
| Embryo | 5 | cis.esQTL.yesChIP | 7 | Liver | 1 | cis.eQTL.yeschip | 7547 | cis.sQTL.yeschip | 9691 |
| Embryo | 6 | cis.esQTL.noChIP | 8 | Liver | 2 | cis.eQTL.nochip | 10519 | cis.sQTL.nochip | 13194 |
| Embryo | 7 | trans.eQTL.yesChIP | 3437 | Liver | 3 | trans.eQTL.yeschip | 2070 | trans.sQTL.yeschip | 11160 |
| Embryo | 8 | trans.eQTL.noChIP | 6838 | Liver | 4 | trans.eQTL.nochip | 5141 | trans.sQTL.nochip | 25035 |
| Embryo | 9 | trans.sQTL.yesChIP | 3080 | Liver | 5 | remaining | 1857218 | remaining | 1823415 |
| Embryo | 10 | trans.sQTL.noChIP | 5246 | Macrophage | 1 | cis.eQTL.yeschip | 605 | cis.sQTL.yeschip | 3761 |
| Embryo | 11 | trans.esQTL.yesChIP | 1181 | Macrophage | 2 | cis.eQTL.nochip | 905 | cis.sQTL.nochip | 5279 |
| Embryo | 12 | trans.esQTL.noChIP | 2150 | Macrophage | 3 | trans.eQTL.yeschip | 2475 | trans.sQTL.yeschip | 13003 |
| Embryo | 13 | remaining | 1857435 | Macrophage | 4 | trans.eQTL.nochip | 4207 | trans.sQTL.nochip | 26495 |
| Hypothalamus | 1 | cis.eQTL.yesChIP | 8516 | Macrophage | 5 | remaining | 1874305 | remaining | 1833958 |
| Hypothalamus | 2 | cis.eQTL.noChIP | 14941 | Mammary | 1 | cis.eQTL.yeschip | 1717 | cis.sQTL.yeschip | 5167 |
| Hypothalamus | 3 | cis.sQTL.yesChIP | 8195 | Mammary | 2 | cis.eQTL.nochip | 2100 | cis.sQTL.nochip | 6928 |
| Hypothalamus | 4 | cis.sQTL.noChIP | 12300 | Mammary | 3 | trans.eQTL.yeschip | 1116 | trans.sQTL.yeschip | 10049 |
| Hypothalamus | 5 | cis.esQTL.yesChIP | 896 | Mammary | 4 | trans.eQTL.nochip | 2090 | trans.sQTL.nochip | 21799 |
| Hypothalamus | 6 | cis.esQTL.noChIP | 974 | Mammary | 5 | remaining | 1875475 | remaining | 1838555 |
| Hypothalamus | 7 | trans.eQTL.yesChIP | 1576 | Milk_cell | 1 | cis.eQTL.yeschip | 11287 | cis.sQTL.yeschip | 42379 |
| Hypothalamus | 8 | trans.eQTL.noChIP | 3662 | Milk_cell | 2 | cis.eQTL.nochip | 21165 | cis.sQTL.nochip | 84032 |
| Hypothalamus | 9 | trans.sQTL.yesChIP | 12308 | Milk_cell | 3 | trans.eQTL.yeschip | 24 | trans.sQTL.yeschip | 1786 |
| Hypothalamus | 10 | trans.sQTL.noChIP | 29335 | Milk_cell | 4 | trans.eQTL.nochip | 75 | trans.sQTL.nochip | 6312 |
| Hypothalamus | 11 | trans.esQTL.yesChIP | 971 | Milk_cell | 5 | remaining | 1849945 | remaining | 1747987 |
| Hypothalamus | 12 | trans.esQTL.noChIP | 2407 | Monocytes | 1 | cis.eQTL.yeschip | 1961 | cis.sQTL.yeschip | 2761 |
| Hypothalamus | 13 | remaining | 1786415 | Monocytes | 2 | cis.eQTL.nochip | 4397 | cis.sQTL.nochip | 4457 |
| Muscle (Cesar et al.) | 1 | cis.eQTL.yesChIP | 1358 | Monocytes | 3 | trans.eQTL.yeschip | 636 | trans.sQTL.yeschip | 11159 |
| Muscle (Cesar et al.) | 2 | cis.eQTL.noChIP | 1801 | Monocytes | 4 | trans.eQTL.nochip | 1481 | trans.sQTL.nochip | 25048 |
| Muscle (Cesar et al.) | 3 | cis.sQTL.yesChIP | 23648 | Monocytes | 5 | remaining | 1874024 | remaining | 1839074 |
| Muscle (Cesar et al.) | 4 | cis.sQTL.noChIP | 39306 | Muscle | 1 | cis.eQTL.yeschip | 5087 | cis.sQTL.yeschip | 7563 |
| Muscle (Cesar et al.) | 5 | cis.esQTL.yesChIP | 971 | Muscle | 2 | cis.eQTL.nochip | 6321 | cis.sQTL.nochip | 10060 |

| | | | |
|---|---|---|---|
| Muscle (Cesar et al.) | 6 | cis.esQTL.noChIP | 1249 |
| Muscle (Cesar et al.) | 7 | trans.eQTL.yesChIP | 338 |
| Muscle (Cesar et al.) | 8 | trans.eQTL.noChIP | 746 |
| Muscle (Cesar et al.) | 9 | trans.sQTL.yesChIP | 8802 |
| Muscle (Cesar et al.) | 10 | trans.sQTL.noChIP | 19020 |
| Muscle (Cesar et al.) | 11 | trans.esQTL.yesChIP | 699 |
| Muscle (Cesar et al.) | 12 | trans.esQTL.noChIP | 1334 |
| Muscle (Cesar et al.) | 13 | remaining | 1783225 |
| Jejunum | 1 | cis.eQTL.yesChIP | 639 |
| Jejunum | 2 | cis.eQTL.noChIP | 916 |
| Jejunum | 3 | cis.sQTL.yesChIP | 3408 |
| Jejunum | 4 | cis.sQTL.noChIP | 5479 |
| Jejunum | 5 | cis.esQTL.yesChIP | 188 |
| Jejunum | 6 | cis.esQTL.noChIP | 302 |
| Jejunum | 7 | trans.eQTL.yesChIP | 386 |
| Jejunum | 8 | trans.eQTL.noChIP | 992 |
| Jejunum | 9 | trans.sQTL.yesChIP | 10921 |
| Jejunum | 10 | trans.sQTL.noChIP | 23556 |
| Jejunum | 11 | trans.esQTL.yesChIP | 499 |
| Jejunum | 12 | trans.esQTL.noChIP | 1033 |
| Jejunum | 13 | remaining | 1834177 |
| Liver | 1 | cis.eQTL.yesChIP | 4945 |
| Liver | 2 | cis.eQTL.noChIP | 7152 |
| Liver | 3 | cis.sQTL.yesChIP | 6924 |
| Liver | 4 | cis.sQTL.noChIP | 9573 |
| Liver | 5 | cis.esQTL.yesChIP | 2602 |
| Liver | 6 | cis.esQTL.noChIP | 3367 |
| Liver | 7 | trans.eQTL.yesChIP | 987 |
| Liver | 8 | trans.eQTL.noChIP | 2552 |
| Liver | 9 | trans.sQTL.yesChIP | 9993 |
| Liver | 10 | trans.sQTL.noChIP | 22312 |
| Liver | 11 | trans.esQTL.yesChIP | 1083 |
| Liver | 12 | trans.esQTL.noChIP | 2589 |

| | | | | | |
|---|---|---|---|---|---|
| Muscle | 3 | trans.eQTL.yeschip | 1869 | trans.sQTL.yeschip | 11463 |
| Muscle | 4 | trans.eQTL.nochip | 4464 | trans.sQTL.nochip | 24886 |
| Muscle | 5 | remaining | 1864755 | remaining | 1828525 |
| Ovary | 1 | cis.eQTL.yeschip | 474 | cis.sQTL.yeschip | 4925 |
| Ovary | 2 | cis.eQTL.nochip | 691 | cis.sQTL.nochip | 7702 |
| Ovary | 3 | trans.eQTL.yeschip | 1621 | trans.sQTL.yeschip | 17073 |
| Ovary | 4 | trans.eQTL.nochip | 3699 | trans.sQTL.nochip | 36543 |
| Ovary | 5 | remaining | 1876014 | remaining | 1816255 |
| Pituitary | 1 | cis.eQTL.yeschip | 7681 | cis.sQTL.yeschip | 21431 |
| Pituitary | 2 | cis.eQTL.nochip | 12449 | cis.sQTL.nochip | 34333 |
| Pituitary | 3 | trans.eQTL.yeschip | 4295 | trans.sQTL.yeschip | 16997 |
| Pituitary | 4 | trans.eQTL.nochip | 10086 | trans.sQTL.nochip | 40204 |
| Pituitary | 5 | remaining | 1847985 | remaining | 1769534 |
| Rumen | 1 | cis.eQTL.yeschip | 1758 | cis.sQTL.yeschip | 4105 |
| Rumen | 2 | cis.eQTL.nochip | 2551 | cis.sQTL.nochip | 5977 |
| Rumen | 3 | trans.eQTL.yeschip | 2055 | trans.sQTL.yeschip | 12828 |
| Rumen | 4 | trans.eQTL.nochip | 4868 | trans.sQTL.nochip | 27737 |
| Rumen | 5 | remaining | 1871265 | remaining | 1831848 |
| Uterus | 1 | cis.eQTL.yeschip | 3179 | cis.sQTL.yeschip | 6305 |
| Uterus | 2 | cis.eQTL.nochip | 4072 | cis.sQTL.nochip | 9404 |
| Uterus | 3 | trans.eQTL.yeschip | 994 | trans.sQTL.yeschip | 10082 |
| Uterus | 4 | trans.eQTL.nochip | 2297 | trans.sQTL.nochip | 24122 |
| Uterus | 5 | remaining | 1871955 | remaining | 1832585 |
| Multi.tissue | 1 | cis.eQTL.yeschip | 106299 | cis.sQTL.yeschip | 226863 |
| Multi.tissue | 2 | cis.eQTL.nochip | 171010 | cis.sQTL.nochip | 528171 |
| Multi.tissue | 3 | trans.eQTL.yeschip | 58790 | trans.sQTL.yeschip | 12884 |
| Multi.tissue | 4 | trans.eQTL.nochip | 218028 | trans.sQTL.nochip | 85791 |
| Multi.tissue | 5 | remaining | 1328374 | remaining | 1028791 |

| | | | |
|---|---|---|---|
| Liver | 13 | remaining | 1808417 |
| Macrophage | 1 | cis.eQTL.yesChIP | 466 |
| Macrophage | 2 | cis.eQTL.noChIP | 715 |
| Macrophage | 3 | cis.sQTL.yesChIP | 3347 |
| Macrophage | 4 | cis.sQTL.noChIP | 4737 |
| Macrophage | 5 | cis.esQTL.yesChIP | 139 |
| Macrophage | 6 | cis.esQTL.noChIP | 190 |
| Macrophage | 7 | trans.eQTL.yesChIP | 725 |
| Macrophage | 8 | trans.eQTL.noChIP | 1064 |
| Macrophage | 9 | trans.sQTL.yesChIP | 11382 |
| Macrophage | 10 | trans.sQTL.noChIP | 23491 |
| Macrophage | 11 | trans.esQTL.yesChIP | 1750 |
| Macrophage | 12 | trans.esQTL.noChIP | 3143 |
| Macrophage | 13 | remaining | 1831347 |
| Mammary | 1 | cis.eQTL.yesChIP | 1018 |
| Mammary | 2 | cis.eQTL.noChIP | 1213 |
| Mammary | 3 | cis.sQTL.yesChIP | 4370 |
| Mammary | 4 | cis.sQTL.noChIP | 5917 |
| Mammary | 5 | cis.esQTL.yesChIP | 699 |
| Mammary | 6 | cis.esQTL.noChIP | 887 |
| Mammary | 7 | trans.eQTL.yesChIP | 541 |
| Mammary | 8 | trans.eQTL.noChIP | 917 |
| Mammary | 9 | trans.sQTL.yesChIP | 9486 |
| Mammary | 10 | trans.sQTL.noChIP | 20658 |
| Mammary | 11 | trans.esQTL.yesChIP | 575 |
| Mammary | 12 | trans.esQTL.noChIP | 1173 |
| Mammary | 13 | remaining | 1835044 |
| Milk_cell | 1 | cis.eQTL.yesChIP | 5044 |
| Milk_cell | 2 | cis.eQTL.noChIP | 10336 |
| Milk_cell | 3 | cis.sQTL.yesChIP | 36131 |
| Milk_cell | 4 | cis.sQTL.noChIP | 73192 |
| Milk_cell | 5 | cis.esQTL.yesChIP | 6243 |
| Milk_cell | 6 | cis.esQTL.noChIP | 10829 |

| | | | |
|---|---|---|---|
| Milk_cell | 7 | trans.eQTL.yesChIP | 19 |
| Milk_cell | 8 | trans.eQTL.noChIP | 61 |
| Milk_cell | 9 | trans.sQTL.yesChIP | 1758 |
| Milk_cell | 10 | trans.sQTL.noChIP | 6252 |
| Milk_cell | 11 | trans.esQTL.yesChIP | 5 |
| Milk_cell | 12 | trans.esQTL.noChIP | 14 |
| Milk_cell | 13 | remaining | 1732614 |
| Monocytes | 1 | cis.eQTL.yesChIP | 1608 |
| Monocytes | 2 | cis.eQTL.noChIP | 3828 |
| Monocytes | 3 | cis.sQTL.yesChIP | 2366 |
| Monocytes | 4 | cis.sQTL.noChIP | 3812 |
| Monocytes | 5 | cis.esQTL.yesChIP | 353 |
| Monocytes | 6 | cis.esQTL.noChIP | 569 |
| Monocytes | 7 | trans.eQTL.yesChIP | 260 |
| Monocytes | 8 | trans.eQTL.noChIP | 504 |
| Monocytes | 9 | trans.sQTL.yesChIP | 10486 |
| Monocytes | 10 | trans.sQTL.noChIP | 23494 |
| Monocytes | 11 | trans.esQTL.yesChIP | 376 |
| Monocytes | 12 | trans.esQTL.noChIP | 977 |
| Monocytes | 13 | remaining | 1833865 |
| Muscle | 1 | cis.eQTL.yesChIP | 3603 |
| Muscle | 2 | cis.eQTL.noChIP | 4435 |
| Muscle | 3 | cis.sQTL.yesChIP | 5924 |
| Muscle | 4 | cis.sQTL.noChIP | 7964 |
| Muscle | 5 | cis.esQTL.yesChIP | 1484 |
| Muscle | 6 | cis.esQTL.noChIP | 1886 |
| Muscle | 7 | trans.eQTL.yesChIP | 871 |
| Muscle | 8 | trans.eQTL.noChIP | 2075 |
| Muscle | 9 | trans.sQTL.yesChIP | 10422 |
| Muscle | 10 | trans.sQTL.noChIP | 22379 |
| Muscle | 11 | trans.esQTL.yesChIP | 998 |
| Muscle | 12 | trans.esQTL.noChIP | 2389 |
| Muscle | 13 | remaining | 1818065 |

| | | | |
|---|---|---|---|
| Ovary | 1 | cis.eQTL.yesChIP | 352 |
| Ovary | 2 | cis.eQTL.noChIP | 553 |
| Ovary | 3 | cis.sQTL.yesChIP | 4577 |
| Ovary | 4 | cis.sQTL.noChIP | 7220 |
| Ovary | 5 | cis.esQTL.yesChIP | 122 |
| Ovary | 6 | cis.esQTL.noChIP | 138 |
| Ovary | 7 | trans.eQTL.yesChIP | 496 |
| Ovary | 8 | trans.eQTL.noChIP | 1199 |
| Ovary | 9 | trans.sQTL.yesChIP | 16066 |
| Ovary | 10 | trans.sQTL.noChIP | 34197 |
| Ovary | 11 | trans.esQTL.yesChIP | 1125 |
| Ovary | 12 | trans.esQTL.noChIP | 2500 |
| Ovary | 13 | remaining | 1813954 |
| Pituitary | 1 | cis.eQTL.yesChIP | 5947 |
| Pituitary | 2 | cis.eQTL.noChIP | 10003 |
| Pituitary | 3 | cis.sQTL.yesChIP | 19018 |
| Pituitary | 4 | cis.sQTL.noChIP | 30695 |
| Pituitary | 5 | cis.esQTL.yesChIP | 1734 |
| Pituitary | 6 | cis.esQTL.noChIP | 2446 |
| Pituitary | 7 | trans.eQTL.yesChIP | 2623 |
| Pituitary | 8 | trans.eQTL.noChIP | 5993 |
| Pituitary | 9 | trans.sQTL.yesChIP | 14810 |
| Pituitary | 10 | trans.sQTL.noChIP | 35391 |
| Pituitary | 11 | trans.esQTL.yesChIP | 1672 |
| Pituitary | 12 | trans.esQTL.noChIP | 4093 |
| Pituitary | 13 | remaining | 1748074 |
| Rumen | 1 | cis.eQTL.yesChIP | 1405 |
| Rumen | 2 | cis.eQTL.noChIP | 2010 |
| Rumen | 3 | cis.sQTL.yesChIP | 3656 |
| Rumen | 4 | cis.sQTL.noChIP | 5287 |
| Rumen | 5 | cis.esQTL.yesChIP | 353 |
| Rumen | 6 | cis.esQTL.noChIP | 541 |
| Rumen | 7 | trans.eQTL.yesChIP | 1077 |

| | | | |
|---|---|---|---|
| Rumen | 8 | trans.eQTL.noChIP | 2464 |
| Rumen | 9 | trans.sQTL.yesChIP | 11791 |
| Rumen | 10 | trans.sQTL.noChIP | 25305 |
| Rumen | 11 | trans.esQTL.yesChIP | 978 |
| Rumen | 12 | trans.esQTL.noChIP | 2404 |
| Rumen | 13 | remaining | 1825225 |
| Uterus | 1 | cis.eQTL.yesChIP | 2198 |
| Uterus | 2 | cis.eQTL.noChIP | 2695 |
| Uterus | 3 | cis.sQTL.yesChIP | 5239 |
| Uterus | 4 | cis.sQTL.noChIP | 7899 |
| Uterus | 5 | cis.esQTL.yesChIP | 981 |
| Uterus | 6 | cis.esQTL.noChIP | 1377 |
| Uterus | 7 | trans.eQTL.yesChIP | 473 |
| Uterus | 8 | trans.eQTL.noChIP | 1021 |
| Uterus | 9 | trans.sQTL.yesChIP | 9545 |
| Uterus | 10 | trans.sQTL.noChIP | 22834 |
| Uterus | 11 | trans.esQTL.yesChIP | 521 |
| Uterus | 12 | trans.esQTL.noChIP | 1276 |
| Uterus | 13 | remaining | 1826436 |
| Multi.tissue | 1 | cis.eQTL.yeschip | 343 |
| Multi.tissue | 2 | cis.eQTL.nochip | 1576 |
| Multi.tissue | 3 | cis.sQTL.yeschip | 67720 |
| Multi.tissue | 4 | cis.sQTL.nochip | 184798 |
| Multi.tissue | 5 | cis.esQTL.yeschip | 105956 |
| Multi.tissue | 6 | cis.esQTL.nochip | 169434 |
| Multi.tissue | 7 | trans.eQTL.yeschip | 53187 |
| Multi.tissue | 8 | trans.eQTL.nochip | 173939 |
| Multi.tissue | 9 | trans.sQTL.yeschip | 7076 |
| Multi.tissue | 10 | trans.sQTL.nochip | 40618 |
| Multi.tissue | 11 | trans.esQTL.yeschip | 5603 |
| Multi.tissue | 12 | trans.esQTL.nochip | 44089 |
| Multi.tissue | 13 | remaining | 1028161 |

**Supplementary Table 3**. Cattle traits analysed in the study with their short name and trait order.Where cow.N is the number of cows for each trait and bull.N is number of bulls for each trait.

| trait full name | short.name | trait.order | cow.N | bull.N |
|---|---|---|---|---|
| protein yield | Prot | tr01 | 76659 | 8097 |
| fat yield | Fat | tr02 | 76659 | 8097 |
| milk yield | Milk | tr03 | 76659 | 8097 |
| protein percentage | ProtP | tr04 | 76659 | 8097 |
| fat percentage | FatP | tr05 | 76659 | 8097 |
| mastitis | Mas | tr06 | 77642 | 8103 |
| somatic cell count | Scc | tr07 | 75429 | 8083 |
| survival | Surv | tr08 | 61056 | 7147 |
| fertility | Fert | tr09 | 56840 | 7254 |
| ease (of birth) | Ease | tr10 | 43970 | 7835 |
| birth size | BSize | tr11 | 43755 | 7827 |
| gestation length | Gl | tr12 | 37214 | 7181 |
| temperament | Temp | tr13 | 37355 | 6966 |
| milking speed | MSpeed | tr14 | 37303 | 6966 |
| likeability | Like | tr15 | 37323 | 6966 |
| stature | Stat | tr16 | 45056 | 7022 |
| chest width | ChestW | tr17 | 45056 | 7022 |
| angularity | Angul | tr18 | 45056 | 7022 |
| bone quality | Bone | tr19 | 45056 | 7022 |
| rear legs set | RSet | tr20 | 45056 | 7022 |
| fore attachment | ForeA | tr21 | 45056 | 7022 |
| rear attachment height | RearAH | tr22 | 45056 | 7022 |
| front teat placement | TeatPF | tr23 | 45056 | 7022 |
| loin strength | Loin | tr24 | 45056 | 7022 |
| udder texture | UdTex | tr25 | 45056 | 7022 |
| central ligament | CentL | tr26 | 45055 | 7022 |
| pin width | PinW | tr27 | 45055 | 7022 |
| foot angle | FootA | tr28 | 45055 | 7022 |
| udder depth | UdDep | tr29 | 45055 | 7022 |
| pin set | PinSet | tr30 | 45054 | 7022 |
| rear attachment width | RearAW | tr31 | 45054 | 7022 |
| muzzle width | MuzW | tr32 | 45054 | 7022 |
| teat length | TeatL | tr33 | 45053 | 7022 |
| body depth | BodyD | tr34 | 45052 | 7022 |
| overall type | OType | tr35 | 44930 | 7022 |
| rear teat placement | TeatPR | tr36 | 44706 | 7022 |
| rear leg view | RLeg | tr37 | 44685 | 7022 |

**Supplementary Table 4.** Summary of the proportion (%) of heritability and trait-associated variants (QTL) in expression quantitative trait loci (eQTL) and RNA splicing QTL (sQTL) when eQTL and sQTL were analysed separately and those e/sQTL were under histone marks ("_Histone"). Within each class, the total number of variants (N class) and their genome proportion (% class) as well as the number of variants with small, medium and large effects averaged across 16 tissues and 37 traits are given. These numbers were used to estimate the observed proportion of heritability explained (O[% $h^2$]) and proportion of QTL in each class (O[% QTL]). The number of variants with in the remaining class (no regulatory evidence) were used to estimate the expected proportion of heritability explained (E[% $h^2$]) and % QTL in each class (E[% QTL]).

| model | tissue | class | N.all | prop.SNP | Small | Medium | Large | O[% $h^2$] (se) | E[% $h^2$] (se) | O[% QTL] (se) | E[% QTL] (se) |
|---|---|---|---|---|---|---|---|---|---|---|---|
| eQTL alone | Single.tissue | cis.eQTL_Histone | 4940 | 0.26 | 110.1(3) | 11.1(0.4) | 0.4(0.0) | 2.59(0.07) | 0.23(0.01) | 4.87(0.18) | 0.001(0.000) |
| | | cis.eQTL | 7579 | 0.40 | 132.1(5) | 12.7(0.5) | 0.4(0.0) | 2.98(0.10) | 0.36(0.02) | 3.58(0.13) | 0.002(0.000) |
| | | trans.eQTL_Histone | 1786 | 0.09 | 71.8(2) | 8.1(0.3) | 0.3(0.0) | 1.79(0.05) | 0.09(0.00) | 7.94(0.41) | 0.000(0.000) |
| | | trans.eQTL | 3958 | 0.21 | 94.4(2) | 11.0(0.4) | 0.3(0.0) | 2.32(0.06) | 0.19(0.01) | 5.00(0.30) | 0.001(0.000) |
| | | remaining | 1864238 | 99.03 | 7649.7(47) | 130.3(2.4) | 1.9(0.1) | 90.33(0.22) | 90.33(0.22) | 0.42(0.00) | 0.417(0.002) |
| | Multi.tissue | cis.eQTL_Histone | 106299 | 5.65 | 691.9(34) | 29.6(3.6) | 0.9(0.2) | 10.68(0.57) | 4.00(0.12) | 0.68(0.03) | 0.027(0.001) |
| | | cis.eQTL | 171010 | 9.08 | 1381.1(76) | 41.4(4.2) | 0.8(0.2) | 18.50(0.97) | 6.44(0.19) | 0.83(0.04) | 0.043(0.002) |
| | | trans.eQTL_Histone | 58790 | 3.12 | 328.8(18) | 19.8(2.5) | 0.4(0.0) | 5.56(0.27) | 2.21(0.06) | 0.59(0.03) | 0.015(0.001) |
| | | trans.eQTL | 218028 | 11.58 | 1234.5(63) | 27.6(3.1) | 0.5(0.0) | 15.25(0.70) | 8.21(0.24) | 0.58(0.03) | 0.054(0.002) |
| | | remaining | 1328374 | 70.56 | 4337.8(157) | 64.3(8.1) | 1.2(0.3) | 50.01(1.44) | 50.01(1.44) | 0.33(0.01) | 0.331(0.012) |
| sQTL alone | Single.tissue | cis.sQTL_Histone | 11932 | 0.63 | 170.3(5) | 14.9(0.5) | 0.5(0.0) | 3.64(0.10) | 0.53(0.02) | 2.66(0.12) | 0.003(0.000) |
| | | cis.sQTL | 19318 | 1.03 | 247.9(11) | 17.9(0.7) | 0.4(0.0) | 4.64(0.16) | 0.86(0.04) | 2.13(0.09) | 0.004(0.000) |
| | | trans.sQTL_Histone | 10843 | 0.58 | 136.9(3) | 15.3(0.5) | 0.4(0.0) | 3.22(0.08) | 0.50(0.01) | 1.67(0.06) | 0.002(0.000) |
| | | trans.sQTL | 24279 | 1.29 | 207.4(4) | 19.5(0.6) | 0.4(0.0) | 4.35(0.09) | 1.13(0.02) | 1.03(0.02) | 0.005(0.000) |
| | | remaining | 1816128 | 96.47 | 7234.9(49) | 113.1(2.2) | 1.8(0.1) | 84.15(0.32) | 84.15(0.32) | 0.40(0.00) | 0.404(0.003) |
| | Multi.tissue | cis.sQTL_Histone | 226863 | 12.05 | 1279.2(42) | 45.0(4.9) | 1.1(0.3) | 18.20(0.78) | 7.45(0.36) | 0.58(0.02) | 0.061(0.003) |
| | | cis.sQTL | 528171 | 28.06 | 3367.6(125) | 56.8(6.7) | 0.9(0.2) | 39.63(1.22) | 17.35(0.83) | 0.65(0.02) | 0.142(0.007) |
| | | trans.sQTL_Histone | 12884 | 0.68 | 126.6(12) | 11.0(1.3) | 0.3(0.0) | 2.60(0.21) | 0.42(0.02) | 1.07(0.09) | 0.003(0.000) |
| | | trans.sQTL | 85791 | 4.56 | 391.9(30) | 15.9(2.0) | 0.4(0.0) | 5.76(0.41) | 2.82(0.14) | 0.48(0.04) | 0.023(0.001) |
| | | remaining | 1028791 | 54.65 | 2798.4(132) | 54.8(10.5) | 1.0(0.2) | 33.80(1.62) | 33.80(1.62) | 0.28(0.01) | 0.277(0.013) |

**Supplementary Table 5**. Annotations of assayed metabolic traits according to Liu et al. 2015 (https://pubmed.ncbi.nlm.nih.gov/28764079/).

| Metabolites | Class | Structure | Species I | Species II | Species III |
|---|---|---|---|---|---|
| PL1 | Phosphatidylserine | PS 34:2 | 16:0/18:2 | 16:1/18:1 | |
| PL2 | Phosphatidylserine | PS 34:1 | 16:0/18:1 | 18:0/16:1 | |
| PL3 | Phosphatidylserine | PS 36:3 | 18:1/18:2 | 18:0/18:3 | |
| PL4 | Phosphatidylserine | PS 36:2 | 18:0/18:2 | 18:1/18:1 | |
| PL5 | Phosphatidylserine | PS 36:1 | 18:0/18:1 | | |
| PL6 | Phosphatidylserine | PS 38:5 | 16:0/22:5 | 18:1/20:4 | 18:0/20:5 |
| PL7 | Phosphatidylserine | PS 38:4 | 18:1/20:3 | 18:0/20:4 | |
| PL8 | Phosphatidylserine | PS 40:5 | 18:0/22:5 | 18:1/22:4 | |
| PL9 | Sphingomyelin | SM 32:1 | d18:1/14:0 | d16:1/16:0 | |
| PL10 | Sphingomyelin | SM 32:0 | d16:0/16:0 | d18:0/14:0 | |
| PL11 | Sphingomyelin | SM 33:1 | d17:1/16:0 | d18:1/15:0 | |
| PL12 | Sphingomyelin | SM 34:1 | d16:1/18:0 | d18:1/16:0 | |
| PL14 | Sphingomyelin | SM 38:1 | d16:1/22:0 | d18:1/20:0 | |
| PL16 | Sphingomyelin | SM 39:1 | d16:1/23:0 | d17:1/22:0 | |
| PL18 | Sphingomyelin | SM 40:2 | d16:1/24:1 | d17:1/23:1 | |
| PL19 | Sphingomyelin | SM 40:1 | d16:1/24:0 | d17:1/23:0 | d18:1/22:0 |
| PL21 | Sphingomyelin | SM 41:2 | d18:1/23:1 | d16:1/25:1 | |
| PL22 | Sphingomyelin | SM 41:1 | d17:1/24:0 | d18:1/23:0 | d16:1/25:0 |
| PL24 | Sphingomyelin | SM 42:2 | d18:1/24:1 | | |
| PL25 | Sphingomyelin | SM 42:1 | d16:1/26:0 | d18:1/24:0 | d19:1/23:0 |
| PL26 | phosphatidylethanolamine | PE 32:1 | 18:1/14:0 | 16:0/16:1 | |
| PL27 | phosphatidylethanolamine | PE 34:2 | 16:1/18:1 | 16:0/18:2 | 17:1/17:1 |
| PL28 | phosphatidylethanolamine | PE 34:1 | 16:0/18:1 | | |
| PL29 | phosphatidylethanolamine | PE 36:4 | 18:1/18:3 | 18:2/18:2 | |
| PL30 | phosphatidylethanolamine | PE 36:3 | 18:1/18:2 | 18:0/18:3 | |
| PL31 | phosphatidylethanolamine | PE 36:2 | 18:1/18:1 | 18:0/18:2 | |
| PL33 | phosphatidylcholine | PC 28:0 | 16:0/12:0 | 14:0/14:0 | |
| PL34 | phosphatidylcholine | PC 30:0 | 16:0/14:0 | | |
| PL35 | phosphatidylcholine | PC 31:0 | 15:0/16:0 | 17:0/14:0 | |
| PL36 | phosphatidylcholine | PC 32:1 | 18:1/14:0 | 16:0/16:1 | |
| PL37 | phosphatidylcholine | PC 32:0 | 16:0/16:0 | 18:0/14:0 | |
| PL38 | phosphatidylcholine | PC 34:3 | 16:0/18:3 | 16:1/18:2 | |
| PL39 | phosphatidylcholine | PC 34:2 | 16:0/18:2 | 16:1/18:1 | |
| PL40 | phosphatidylcholine | PC 34:1 | 16:0/18:1 | | |
| PL42 | phosphatidylcholine | PC 36:4 | 18:1/18:3 | 16:0/20:4 | 18:2/18:2 |
| PL43 | phosphatidylcholine | PC 36:3 | 18:1/18:2 | 18:0/18:3 | 16:0/20:3 |
| PL44 | phosphatidylcholine | PC 36:2 | 18:1/18:1 | 18:0/18:2 | |
| PL46 | phosphatidylinositol | PI 34:1 | 16:0/18:1 | 18:0/16:1 | |

| | | | | | |
|---|---|---|---|---|---|
| PL47 | phosphatidylinositol | PI 36:2 | 18:1/18:1 | 18:0/18:2 | |
| PL48 | phosphatidylinositol | PI 36:1 | 18:0/18:1 | | |
| PL49 | phosphatidylinositol | PI 38:5 | 18:0/20:5 | 18:1/20:4 | 16:0/22:5 |
| PL50 | phosphatidylinositol | PI 38:4 | 18:1/20:3 | 18:0/20:4 | |
| PL51 | phosphatidylinositol | PI 38:3 | 18:0/20:3 | 18:1/20:2 | |
| PL52 | Lysophosphatidylcholine | LPC 16:0 | 16:0 | | |
| PL53 | Lysophosphatidylcholine | LPC 18:3 | 18:3 | | |
| PL54 | Lysophosphatidylcholine | LPC 18:2 | 18:2 | | |
| PL55 | Lysophosphatidylcholine | LPC 18:1 | 18:1 | | |
| PL56 | Lysophosphatidylcholine | LPC 18:0 | 18:0 | | |
| PL57 | lactosylceramide | LacCer 34:1 | d18:1/16:0 | | |
| PL58 | lactosylceramide | LacCer 38:1 | d16:1/22:0 | | |
| PL59 | lactosylceramide | LacCer 38:0 | d16:0/22:0 | d22:0/16:0 | |
| PL60 | lactosylceramide | LacCer 39:1 | d17:1/22:0 | d18:1/21:0 | d16:1/23:0 |
| PL61 | lactosylceramide | LacCer 39:0 | d16:0/23:0 | | |
| PL62 | lactosylceramide | LacCer 40:1 | d16:1/24:0 | d17:1/23:0 | d18:1/22:0 |
| PL64 | lactosylceramide | LacCer 41:1 | d18:1/23:0 | d18:0/23:1 | |
| PL66 | lactosylceramide | LacCer 42:1 | d18:1/24:0 | | |
| PL67 | Glucosylceramide | GluCer 34:1 | d18:1/16:0 | | |
| PL69 | Glucosylceramide | GluCer 40:1 | d18:1/22:0 | d16:1/24:0 | d17:1/23:0 |
| PL70 | Glucosylceramide | GluCer 42:1 | d18:1/24:0 | | |
